# Direct visualization of MCM helicase activation and replisome coupling in situ

**DOI:** 10.64898/2026.08.21.746258

**Authors:** Oraya J. Zinder, Jonas Zähringer, Hana Polasek-Sedlackova, Kannanganattu V. Prasanth, Taekjip Ha, Supriya G. Prasanth

## Abstract

Deciphering the spatial organization of molecular machines that copy the genome remains a fundamental challenge in biology. Essential for eukaryotic DNA replication, Mini-Chromosome Maintenance (MCM2-7) helicases are loaded during G1 as double hexamers (DHs) to license replication origins. Upon activation in S phase, each DH is thought to split into two single hexamers (SHs) that form the active CMG helicases and travel bidirectionally. However, the field has long been divided: biochemical and structural studies define CMG helicases as autonomous, independent motors, while genomic and cellular imaging assays suggest sister replisomes remain physically coupled within replication factories. Here, we use MINFLUX nanoscopy to localize individual MCM complexes down to nanometer precision in situ, directly resolving DHs in human cells and capturing their separation into SHs upon origin firing. We find that the resulting sister replisomes do not diffuse apart: they remain coupled at a characteristic distance of ∼40 nm throughout S phase. Depletion experiments identify two distinct contributions to this coupling: local, protein-mediated tethering by the AND1 scaffold, and higher-order spatial confinement dependent on cohesin, which is dispensable for MCM loading in G1 but required to maintain coupling in S phase. By linking the nanometer-scale architecture of the replisome to the genome-wide topology of replication fountains, these findings provide direct spatial evidence that sister forks are coupled during DNA synthesis and define the molecular forces that organize replisomes within their native nuclear context.

---

Initiation of DNA replication requires the assembly of a multi-protein prereplicative complex on replication origins during G1 phase^1^. These origins are licensed by the loading of the minichromosome maintenance (MCM) proteins, which assemble in an inactive, head-to-head double hexameric (DH) configuration^2,3^. The binding of the Origin Recognition Complex (ORC) to origins, followed by CDC6 association, drives the recruitment of CDT1-MCM2-7^4–7^. Upon loading the first MCM ring, ORC-CDC6 rebinds to recruit a second CDT1-MCM2-7 complex^8,9^. As cells transition into S-phase, this MCM DH is converted into two single hexamers (SHs), each bound to CDC45 and GINS, forming the active CDC45-MCM2-7-GINS (CMG) helicase^10–13^, which is key to origin activation.

Despite a detailed molecular framework for replication initiation, fundamental questions regarding the quantitative regulation and spatial organization of origin activation remain unresolved. Addressing these questions has been hindered by the lack of approaches capable of resolving origin activation at the level of individual replication forks in their native cellular environment. As a result, how licensed origins are selected and activated in vivo remains largely unknown. MCM complexes are loaded onto chromatin in substantial excess, estimated at 3- to 10-fold more than the number of origins that actually fire during an unperturbed S-phase^14–17^. The majority of these excess MCMs occupy dormant origins: licensed sites that show no detectable initiation activity under normal conditions but can be activated as a reserve when replication forks stall or collapse under replicative stress^18^. ATR kinase plays a central role in regulating this reserve, globally suppressing dormant origin firing in response to stalled forks^18^. The limited colocalization of MCM with markers of active replication, including PCNA and EdU-labeled nascent DNA, supports the idea that only a minority of loaded MCMs are engaged as active helicases at any given time^19^. While the internal molecular architecture of the individual eukaryotic replisome is well defined^20^, the higher-order spatial relationship between sister replisomes remains a subject of intense debate. Live-cell imaging and recent genomic proximity assays suggest that sister replication forks remain physically coupled^21,22^. However, this macroscopic factory model conflicts with biophysical and structural evidence showing that sister CMG helicases physically uncouple and operate as autonomous motors^23–28^. It remains unclear whether sister replisomes are rigidly tethered, completely independent, or maintained in loose spatial proximity below the diffraction limit of standard imaging.

The trimeric scaffold AND1 (Ctf4 in yeast) has been proposed to tether sister CMGs at the molecular level^29^, while the cohesin complex, enriched at replication origins and essential for sister chromatid cohesion, has been implicated in organizing groups of origins within higher-order chromatin domains^30–32^. However, the physical distances at which sister replisomes are coupled and the distinct contributions of protein-mediated tethers, chromatin architecture, and topological constraints have not been directly measured.

To overcome this barrier, we employed MINFLUX nanoscopy, which enables single-molecule localization with a spatial precision of ∼1-5 nm^33^, allowing us to resolve the molecular architecture of the replisome within its native cellular environment. By leveraging the molecular-scale resolution of MINFLUX, we directly resolve the assembly states of individual MCM complexes in situ, defining their spatial organization within replication factories and revealing the molecular mechanisms that couple sister replisomes during bidirectional replication.

## RESULTS

### MCM double hexamers are directly visualized in human cells

The heterohexameric MCM2-7 complex is loaded during G1 as a double hexamer (DH). To test whether DHs can be visualized *in situ*, we used a human osteosarcoma cell line (U2OS) expressing MCM2-Halo at the endogenous locus^34^. Western blot analysis confirmed that all alleles of MCM2 were tagged (Fig. 1a), and HaloTrap immunoprecipitation demonstrated MCM2-Halo’s ability to pull down all the components of the MCM hexamer (Fig. 1b; additional validation in^35^). MCM2-Halo labeled with Abberior FLUX 640-Halo ligand showed colocalization with MCM3 immunofluorescence, with both matching the established MCM spatial patterns (Fig. 1c). Chromatin-bound MCM3 was detected in both G1 and S phases and exhibited distinct spatiotemporal patterning, as confirmed by co-labeling with YFP-ORC1^36^ and PCNA, respectively (Extended Fig. 1a). MCM2-Halo exhibited similar patterns: homogenous labeling in G1 (Fig. 1c, Extended Fig. 1b) and distinct perinucleolar and perinuclear labeling in EdU-positive S-phase cells (Extended Fig. 1b), consistent with the known spatiotemporal progression of MCM through the cell cycle^37^.

**Figure 1.**
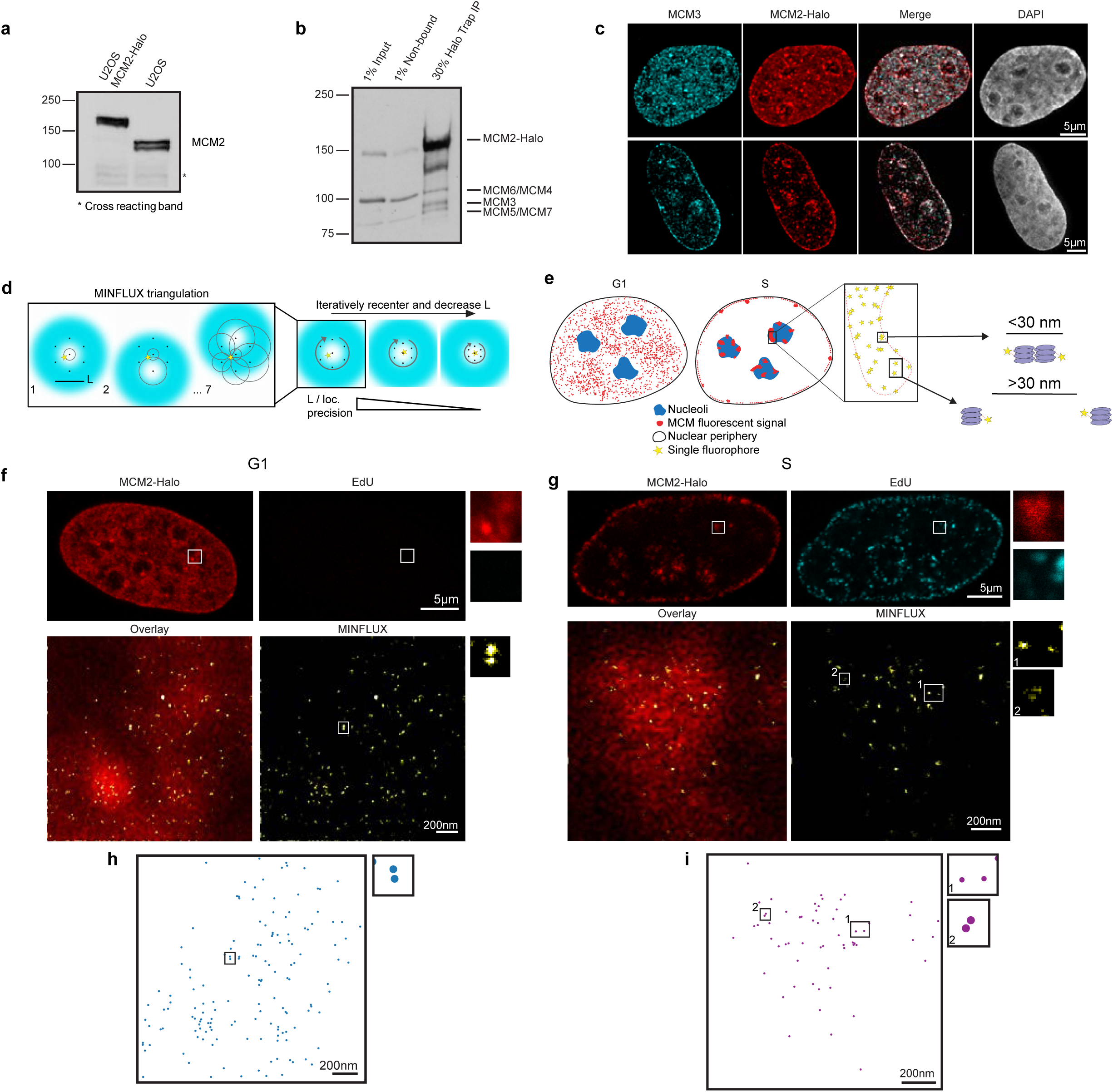
Visualizing MCM double hexamers in vivo. **a.** Western blot analysis of MCM2 in U2OS cells expressing endogenously Halo-tagged MCM2. **b.** HaloTag pull-down immunoprecipitation (Halo Trap) from MCM2-Halo cells. **c**. MCM3 (cyan) immunostaining in MCM2-Halo (red) cells. Top: early S phase. Bottom: mid-late S phase. **c.** Schematic of MINFLUX localization technique, left showing MINFLUX triangulation method, right showing decrease in L with iterative recentering. As L decreases, localization (loc.) precision becomes more precise. **e.** Schematic of MINFLUX quantification and definition of nearest neighbor parameters (<30 nm = DH; >30 nm = SH or split DH). **f**. Representative MINFLUX confocal scan of G1 cell Top left: MCM2-Halo (red); Top right: 10-minute pulse EdU negative cell (scale bar 5 µm); inset: chosen ROI for MINFLUX; Bottom left: MINFLUX localizations overlay with confocal ROI; Bottom right: MINFLUX localizations (yellow, scale bar 200 nm); inset: example DH ∼28 nm. (White inset in top row: ROI for MINFLUX imaging. Inset in bottom row: example DH, ∼28 nm). **g**. Representative MINFLUX confocal scan of S phase cell Top left: MCM2-Halo (red); Top right: 10-minute pulse EdU positive cell (cyan, scale bar 5 µm); inset: chosen ROI for MINFLUX; Bottom left: MINFLUX localizations overlay with confocal ROI; Bottom right: MINFLUX localizations (yellow, scale bar 200 nm); inset: 1. example SH, ∼48 nm 2. example DH, ∼19 nm. (inset in top row: ROI for MINFLUX imaging. Inset in bottom row: 1. example SH, ∼48 nm 2. example DH, ∼19 nm). **h-i**. Representative G1 and S phase dot plots depicting confirmed MCM molecules from MINFLUX localizations. For ease of comparison, (∼160 nm, G1; ∼115 nm S) around the edges of the data were cropped in, full dot plots can be found in Extended Fig. 1f-g. (**h**. inset: representative DH, ∼28 nm; **i**. insets: 1. example SH, ∼48 nm 2. example DH, ∼19 nm).

Next, we used 2D MINFLUX imaging to localize MCM2 at nanometer resolution. MINFLUX uses a donut-shaped excitation profile to achieve single fluorophore localization with precisions down to 1 nm^33^ through successive triangulations close to the excitation minima and progressive refinement (see schematic, Fig. 1d)^33^. If both MCM2 molecules in a DH are detected, we expect the 3D distance between the fluorophores on the C-termini to be approximately 23 nm based on available structural models^38,39^, yielding an average 2D projection of DH distance below 30 nm (Fig. 1e). After confocal imaging, we selected a nuclear region for MINFLUX imaging (Fig. 1f-g: inset in top row). With MINFLUX, individual MCM2 molecules can be resolved within each region of interest (ROI) (Fig. 1f and Extended Fig. 1c). Visual inspection of the raw images readily identified DH-like structures (less than 30 nm spacing); for example, a pair of MCM2 spots separated by 28 nm in G1 (Fig. 1f) and by 19 nm in S phase (Fig. 1g and Extended Fig. 1d).

Following stringent filtering of raw localizations to remove noise and poor localization signals (see Methods), we used density-based spatial clustering of applications with noise (DBSCAN) to identify dense regions of localizations. We then applied K-means clustering, evaluated using the silhouette criterion, to separate individual MCM2 molecules within those DBSCAN clusters (Extended Fig. 1e). The output of our analysis is a localization map, with each point representing the center of mass of a detected MCM2 molecule (Fig. 1h, Extended Fig. 1f-g). Consistent with the raw localizations, we frequently observed doublets, indicating a prominent DH population in G1 (Fig. 1h, Extended Fig. 1f) and in S phase (Fig. 1i, Extended Fig. 1g).

To confirm that the DH population is not undercounted due to incomplete labeling, we performed a Halo-tag ligand titration in the Halo-MCM2 line (500 pM - 2 µM) and established that the fraction of nearest-neighbor (NN) distances < 30 nm saturated between 500 nM and 1 µM (Extended Fig. 2a-b; note merging of yellow & green spots in b); all subsequent experiments were performed at these saturating concentrations. Importantly, the lack of < 30 nm pairs at low labeling yields confirms that the observed DHs represent genuinely coupled MCM complexes rather than MINFLUX localization artifacts.

To assess the statistical significance of the DH population identified from our nearest neighbor analysis, we modeled a random baseline for each imaged region by computationally distributing the total number of molecules across the ROI as independent monomers (Methods and Extended Fig. 2c-d). When compared against this simulated monomeric distribution, the observed DH frequency was significantly enriched, confirming the formation of DH complexes (Fig. 2a-b). Similarly, S-phase populations showed a significant enrichment of DHs over the simulated distribution (Fig. 2c), indicating that these complexes are maintained across both cell cycle stages (Fig. 2d).

**Figure 2.**
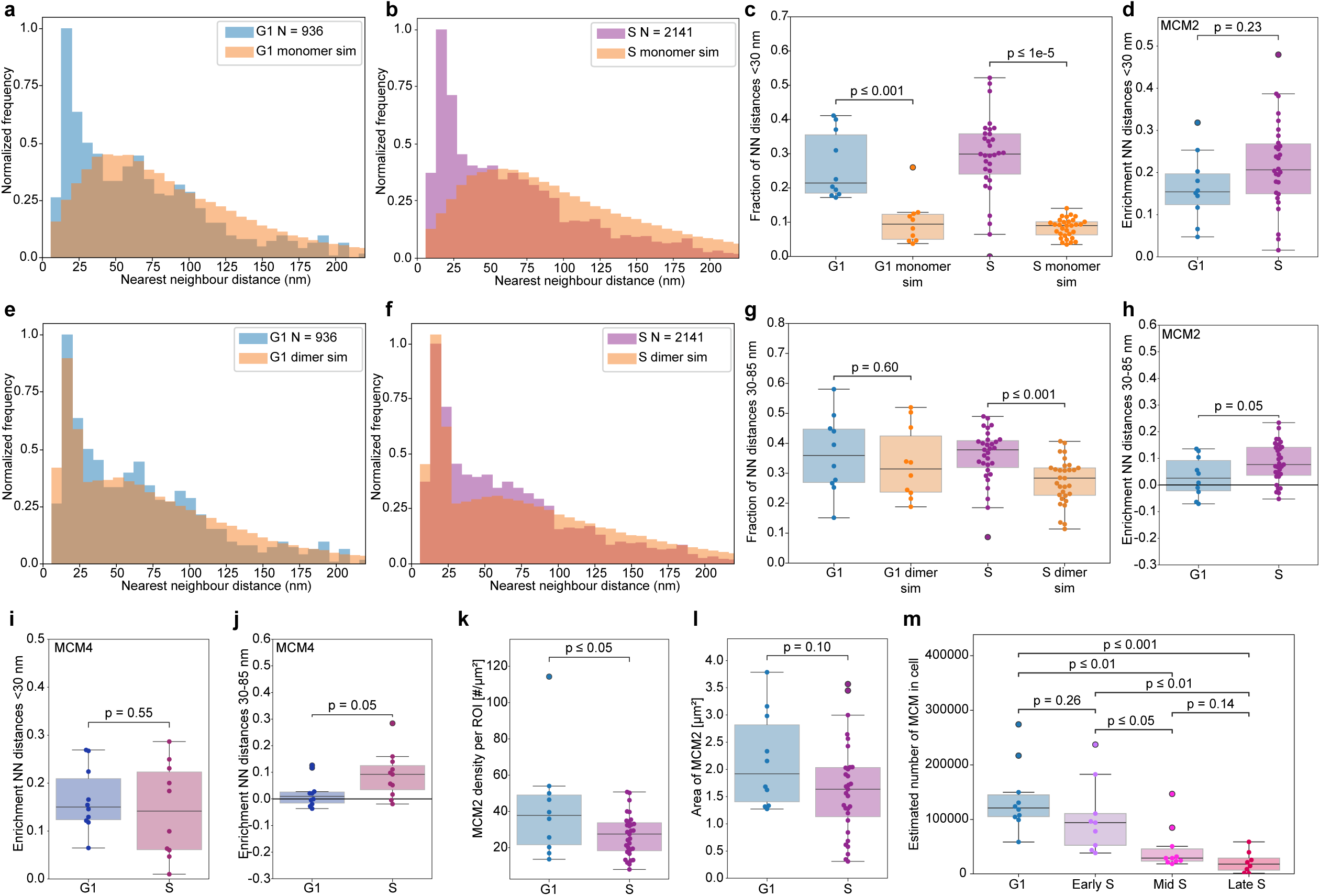
Double hexamer transition into single hexamer of MCMs in S phase. **a-b**. Histograms of nearest neighbor distance (nm) between MCM2 molecules for compiled G1 and S-phase data, respectively, compared to monomeric simulation distributions. **c**. Box plot depicting chromatin-bound DH population for G1 (p ≤ 0.001) and in S (p ≤ 1e-5) compared to monomer simulation distribution. **d**. Box plot showing chromatin-bound DH population from G1 to S for MCM2-Halo (p = 0.23). Note that DH population is not significantly changed from G1 to S for MCM2-Halo. **e-f.** Histograms of nearest-neighbor distances (nm) between MCM2 molecules for compiled G1 and S-phase data, respectively, compared with simulated dimeric distributions. **g**. Box plot depicting chromatin-bound SH population for G1 (p=0.60, non-significant), and in S (p ≤ 0.001) compared to dimer simulation distributions. **h**. Box plot showing enrichment of chromatin-bound SH population in S compared to G1 (p=0.05). Note that SH population is significantly increased from G1 to S for MCM2-Halo. **i**. Box plot showing chromatin-bound DH population from G1 to S (p = 0.55). Note that DH population is not significantly changed from G1 to S for MCM4-Halo, consistent with our observations for MCM2-Halo. **j**. Box plot showing enrichment of chromatin-bound SH population in S compared to G1 for MCM4-Halo (p=0.05). Note that SH population is significantly increased from G1 to S for MCM4-Halo, similar to our observations for MCM2-Halo. **k**. Box plot shows the number of chromatin-bound MCM (#) per micron squared significantly decreasing from G1 to S (p ≤ 0.05). **l**. Box plot showing the area of chromatin-bound MCM from G1 to S (p = 0.10). **m.** Box plot showing the estimated number of chromatin-bound MCM within the cell. Note the decrease from G1 to early to mid to late S-phase. G1 to early S p = 0.26, G1 to mid S p ≤ 0.01, G1 to late S p ≤ 0.001, early S to mid S p ≤ 0.05, early S to late S p ≤ 0.01, mid S to late S p = 0.14. Statistical analyses are included in Supplementary table 1.

### Origin firing converts MCM double hexamers into separated single hexamers

To investigate the conversion of DHs into active single hexamers (SHs) during S-phase, we analyzed populations with NN distances >30 nm. To ensure these greater distances were not simply chance proximities of dense clusters, we developed a mixed spatial simulation incorporating both independent monomers and dimers, with their relative populations tuned to match our overall NN distance distributions. In G1 cells, the >30 nm population showed no significant enrichment over this dimeric simulation distribution (Fig. 2e, g). In contrast, S-phase cells exhibited a distinct and significant enrichment of NN distances specifically between 30 and 85 nm that could not be explained by the simulation (Fig. 2f-h). We propose that this specific 30-85 nm population represents the conversion and physical separation of DHs into SHs during origin firing.

We observed identical dynamics of DH formation and conversion to SHs using a different subunit, MCM4, Halo-tagged at its endogenous locus (Fig. 2i-j, Extended Fig. 3a-f). However, in MCM4, the contrast between DH and SH populations is more distinct. We speculate that an almost four times longer linker on MCM4-Halo (peer review file^19^, 29 aa in MCM2-Halo; 104 aa in MCM4-Halo) likely increases HaloTag’s conformational freedom around the MCM complex, allowing for more distinct distributions of DH and SH (Extended Fig 3a-f).

As replication forks advance, they encounter and unload unfired MCM complexes through mechanisms that remain incompletely understood^13,37,40,41^ ^42^. Consistent with this, our imaging revealed a marked, progressive reduction in chromatin-bound MCM from G1 through S-phase (Fig. 2k). To characterize MCM dynamics across S-phase stages, we utilized EdU co-staining to sub-categorize S-phase into early, mid, and late stages ^37^. Notably, the characteristic MCM pattern precedes the appearance of PCNA- and EdU-positive replication patterns^37^, and MCM is unloaded from chromatin before S-phase concludes (Extended Fig. 4a-e). While the total MCM population (Extended Fig. 4f) and the overall area of MCM foci (Fig. 2l, Extended Fig. 4g) decreased progressively, reflecting a shift from euchromatic to compact heterochromatic domains, the DH fraction remained significantly enriched over the simulation distribution throughout the entirety of S-phase (Extended Fig. 5a-c, g). Concurrently, the enriched SH fraction increased as S-phase progressed (Extended Fig. 4h, Extended Fig. 5d-f, h), likely mirroring the continuous firing of origins.

Next, we leveraged our single-molecule localization data to estimate the absolute number of chromatin-bound MCM complexes. By calibrating the confocal fluorescence signal against the number of MINFLUX-detected MCMs and extrapolating across the full nuclear volume (∼750 µm³ for U2OS), we corrected for a ∼600 nm MINFLUX imaging depth^43^, nuclear Halo-tag labeling efficiency^44^, fluorophore dark states^45^, and baseline detection efficiency^46^. This quantification yielded approximate counts of 120,000 - 160,000 chromatin-bound MCM molecules in G1, dropping to 80,000 - 130,000 in early S-phase, 30,000 - 60,000 in mid S-phase, and 10,000 - 30,000 in late S-phase (Fig. 2m).

### A fraction of loaded MCMs is active within replication factories

In mammalian cells, DNA replication occurs within discrete, spatially clustered replication factories^47–49^. Conventional imaging has historically shown that the bulk of loaded MCM proteins does not colocalize with these active replication sites. This lack of overlap led to the “MCM paradox,” which posits that active MCMs might function at a distance from the replication fork^17^. However, an estimated 3- to 10-fold excess of MCMs is loaded onto chromatin to protect against replicative stress^14–16^. Although a recent study detected endogenous MCM proteins within replicating regions^19^, it lacked the spatial resolution to distinguish inactive DHs from activated CMG helicases within individual replication factories. Consequently, the local organization of active and inactive MCM populations during DNA replication remained unresolved.

To investigate MCM organization at active origins, we overlaid MINFLUX MCM2-Halo localizations with confocal images of EdU-positive replication foci. Strikingly, the vast majority of the MCM signal resided outside the EdU-positive regions (Fig. 3a). However, by setting a stringent EdU confocal signal threshold, we isolated a sparse but distinct population of MCMs directly within the EdU-positive regions, which we infer to be active replication factories (Fig. 3b).

**Figure 3.**
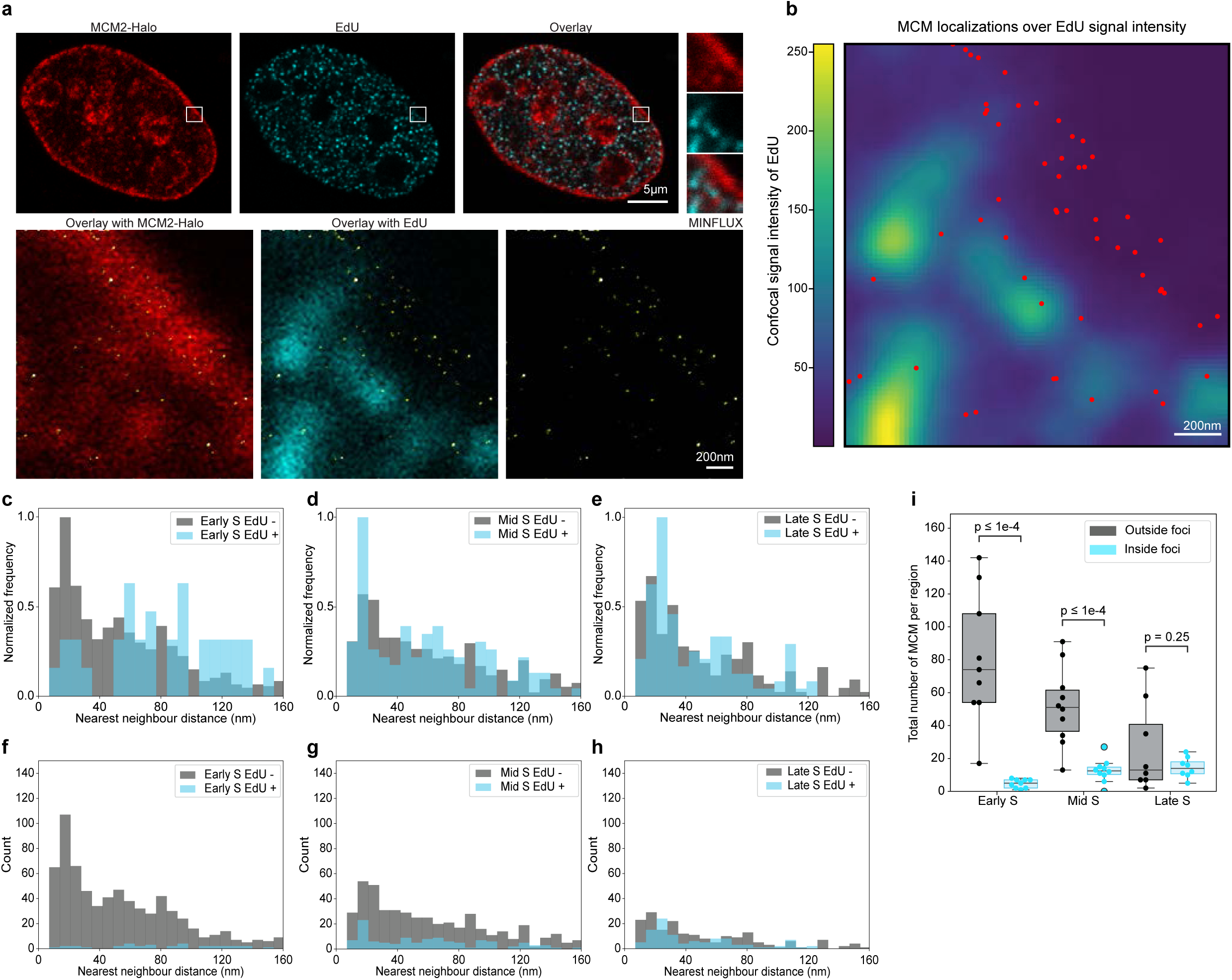
MCM single hexamers show higher overlap with EdU signal. **a.** Representative MINFLUX confocal scan of S phase cell. Top left: MCM2-Halo (red); Top middle: 10-minute pulse EdU positive cell (cyan); Top right: Merge of MCM2-Halo and EdU signal (scale bar 5 µm) and inset: ROI chosen for MINFLUX; Bottom left: MINFLUX localizations overlay with MCM2-Halo confocal ROI; Bottom middle: MINFLUX localizations overlay with EdU confocal ROI; Bottom right: MINFLUX localizations (yellow, scale bar 200 nm). (insets in top row: ROI for MINFLUX imaging). **b**. Dot plot depicting confirmed MCM molecules (red) overlay with EdU signal (green). **c-e**. Normalized density histograms for nearest neighbor distance (nm). Note the rightward shift of MCM molecules that overlap with EdU signal intensity >60 in early S (left), mid S (middle) and late S (right). Note that in early S phase, the gray bars show an increased frequency of nearest-neighbor distances below 30 nm, whereas the blue bars shift rightward. **f-h**. Absolute number of MCM molecules that overlap with EdU signal intensity >60 in early S (left), mid S (middle) and late S (right) plotted as a histogram. **i**. Box plot showing chromatin-bound MCM in EdU positive region during S-phase. Note a significant decrease during early and mid S phase, and a nonsignificant decrease in the number of MCM within EdU-positive regions during late S phase. Statistical analyses are included in Supplementary table 2.

Applying our nearest-neighbor analysis, we found that MCMs in EdU-negative regions were predominantly in the inactive DH configuration (<30 nm) (Fig. 3c-e). Conversely, within EdU-positive regions, there was a dramatic enrichment of the SH configuration (30-85 nm), representing active, spatially separated helicases (Fig. 3c). Across the entirety of S-phase, these SHs constituted the majority of the MCMs within EdU-positive domains (Fig. 3c-e), whereas the absolute number of MCMs inside these domains remained low compared to the bulk chromatin (Fig. 3f-i). We estimate 4-17 MCM molecules per individual replication factory, as defined by the EdU-positive signal, indicating the presence of 2-8 active replicons per factory (Fig 3i). These measurements establish MINFLUX as a powerful tool for directly capturing origin activation in situ and resolving long-standing spatial ambiguities in DNA replication.

### ATR inhibition mobilizes dormant MCM reserves

In metazoans, the vast majority of licensed origins remain dormant during unperturbed S-phase, serving as a backup reserve that fires primarily in response to replicative stress^18^. ATR kinase plays a central role in this regulation: when forks stall, while local origins fire to rescue replication, ATR suppresses global dormant origin firing to limit the number of active replisomes. To directly visualize dormant origin activation, we combined mild replication stress (HU; 0.2 mM) with ATR inhibition (ATRi; NU6027), which abrogates this checkpoint and permits widespread firing of dormant origins, a response confirmed by western blot (Extended Fig. 6a).

MINFLUX imaging of ATRi+HU-treated cells showed that the inactive DH population (NN < 30 nm) remained comparable to that of G1 cells (Fig. 4c-d, Extended Fig. 6b-c). This suggests that only a small fraction of the dormant pool is converted. In contrast, the treated cells exhibited a sharply defined nearest-neighbor peak concentrated specifically within the 30-60 nm range (Fig. 4c). We interpret this as a significant enrichment of the SH fraction, indicating enhanced origin firing (Fig. 4e, Extended Fig. 6d).

**Figure 4.**
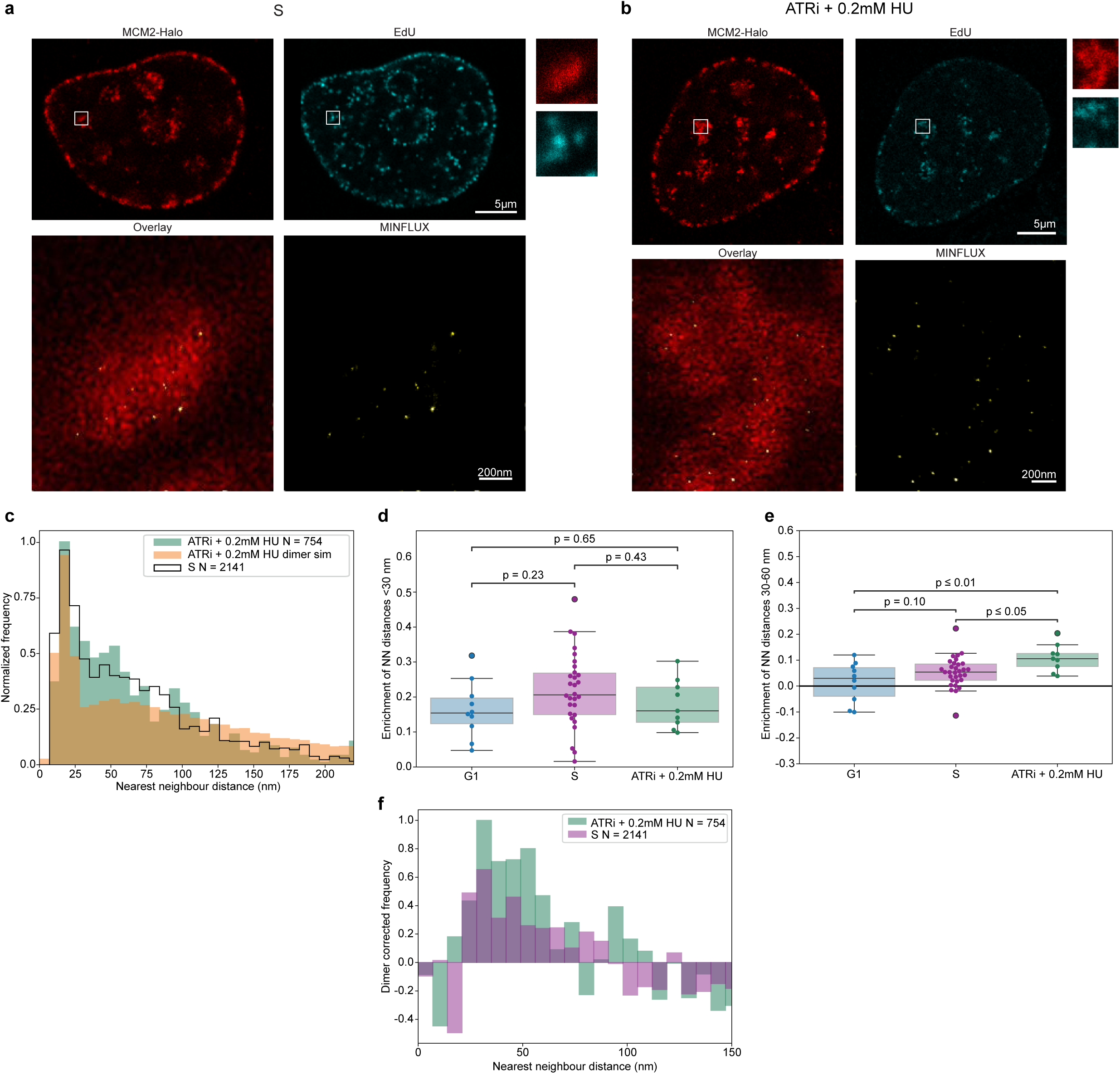
Dormant origin activation increases chromatin bound MCM. **a.** Representative MINFLUX confocal scan of S-phase cell. Top left: MCM2-Halo (red); Top left: 10-minute pulse EdU positive cell (cyan, scale bar 5 µm); inset: chosen ROI for MINFLUX; Bottom left: MINFLUX localizations overlay with confocal ROI; Bottom right: MINFLUX localizations (yellow, scale bar 200 nm). (inset in top row: ROI for MINFLUX imaging.) **b**. Representative MINFLUX confocal scan of ATRi+0.2 mM HU treated S phase cell. Top left: MCM2-Halo (red); Top left: 30-minute pulse EdU positive cell (cyan, scale bar 5 µm); inset: chosen ROI for MINFLUX; Bottom left: MINFLUX localizations overlay with confocal ROI; Bottom right: MINFLUX localizations (yellow, scale bar 200 nm). (inset in top row: ROI for MINFLUX imaging.) **c**. Histogram of nearest neighbor distance (nm) between MCM2 molecules for compiled ATRi+0.2 mM HU-treated S-phase cells (green) compared to dimeric simulation distribution (orange) and untreated S-phase population (black line). **d**. Box plot showing chromatin-bound DH population from G1, S and ATRi+0.2 mM HU treated. A notable decrease in DH population from S to ATRi+0.2 mM HU treated cells is seen. **e**. Box plot showing chromatin-bound SH population from G1 to S and ATRi+0.2 mM HU treated. Note the significant enrichment of SH population from G1 to S and to ATRi+0.2 mM HU and ATRi+0.2 mM HU treated compared to S-phase cells. **f.** Dimer corrected frequency histogram of S and ATRi+0.2 mM HU. Note the increase of SH population 30-60 nm. Statistical analyses are included in Supplementary table 3.

Because this ATRi-HU-induced peak is narrow, a feature made clearer through the dimer-corrected frequency (Fig. 4f, Extended Fig. 6g-h), we restricted our SH population analysis to this 30-60 nm window. We suspect that this highly constrained distance reflects a state in which sister helicases remain physically coupled via protein tethering. Including broader distances (60-85 nm) diminished the observed difference (Extended Fig. 6e-f), likely because this wide range captures uncoupled MCM complexes. This suggests that ATR inhibition under stress preferentially traps a spatially restricted population of SHs rather than causing a broad redistribution, reflecting altered replisome dynamics and slowed fork progression^50^. We also observed a decrease in overall MCM density per ROI (Extended Fig. 6i) alongside an increase in the spatial area occupied by MCMs (Extended Fig. 6j), indicating that stress-induced firing disperses the MCM complexes within replication foci.

### AND1 and cohesin maintain sister replisome coupling through distinct mechanisms

Next, we examined whether sister replisomes remain physically coupled during active DNA replication. Cryo-EM studies in budding yeast have demonstrated that two CMG helicases and a single Pol α-primase are held together by the trimeric scaffold protein Ctf4 (AND1 in higher eukaryotes)^29^. To test whether AND1 mediates this coupling in human cells, we depleted AND1 using siRNA (Fig. 5a, Extended Fig. 7a). Importantly, the non-targeting control siGL3 showed results consistent with the S-phase control (Extended Fig. 7d-j). AND1 depletion did not significantly alter total MCM protein levels, cell cycle distribution, or EdU incorporation (Fig. 5a, Extended Fig. 7a, c), confirming that global replication was not broadly disrupted. AND1 depletion caused a modest decrease in the DH population without significantly altering the overall SH fraction (Fig. 5b, d-e, Extended Fig. 8a, c-d); the basis for this DH reduction is not yet fully understood and may reflect secondary effects on replication dynamics.

**Figure 5.**
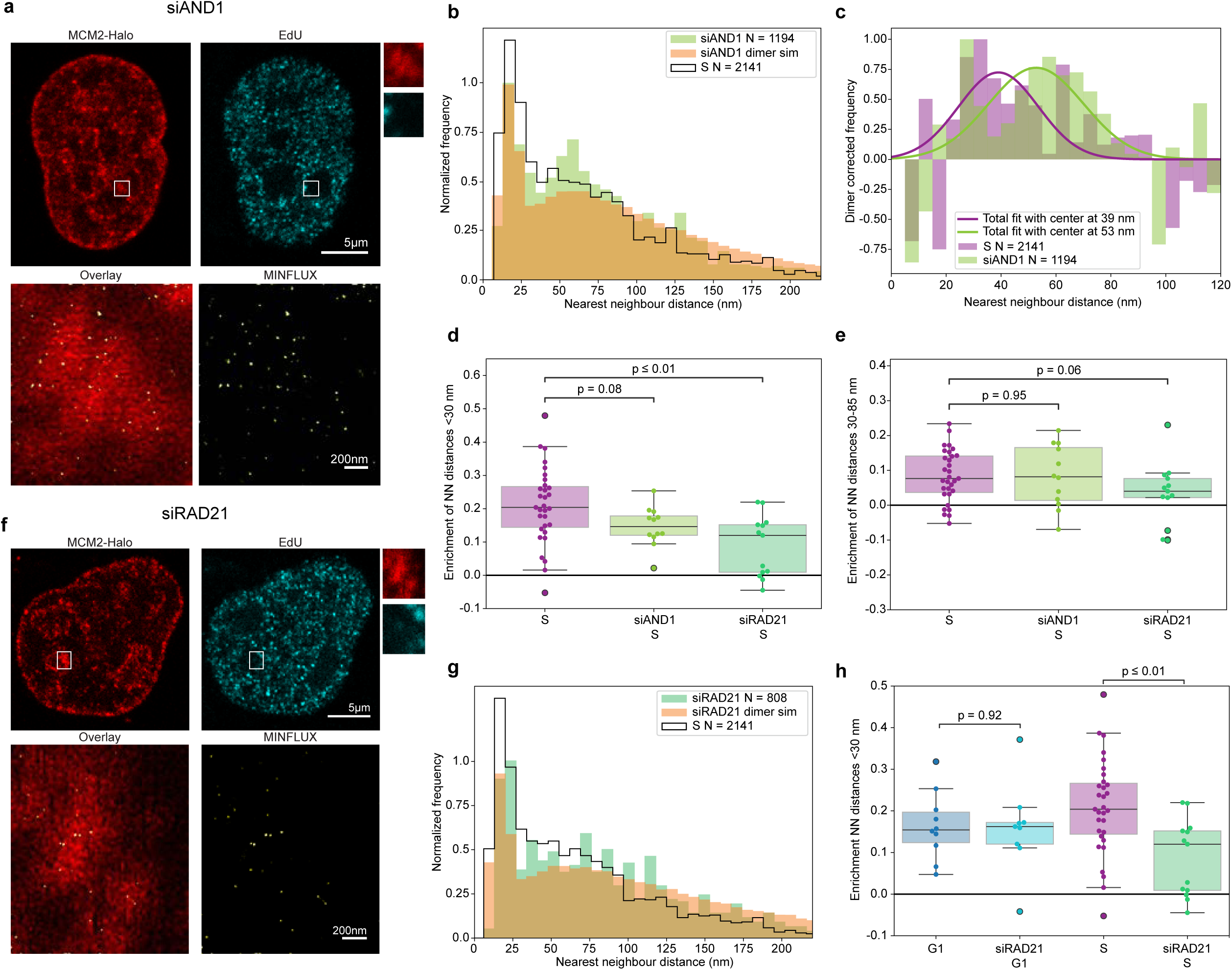
Replisome coupling promote the nanoscale organization of chromatin-bound MCM2 during S phase. **a.** Representative MINFLUX images of siAND1-treated cells respectively. MINFLUX confocal scan of S phase cell. Top left: MCM2-Halo (red); Top right: 10-minute pulse EdU positive cell (cyan, scale bar 5 µm); inset: chosen ROI for MINFLUX; Bottom left: MINFLUX localizations overlay with confocal ROI; Bottom right: MINFLUX localizations (yellow, scale bar 200 nm). (inset in top row: ROI for MINFLUX imaging.) **b.** Histogram of nearest-neighbor distances (nm) between MCM2 molecules for compiled S-phase data from siAND1-treated cells (green) with the simulated dimeric distribution (orange) and untreated S-phase population (black line). **c.** Dimer corrected frequency histogram of S-phase (total fit center at 39 nm), and siAND1 (total fit center at 53 nm). Note the 14 nm shift in gaussian fit center from S-phase to siAND1-treated S phase cells (p = 0.04). **d.** Box plot showing enrichment of chromatin-bound DH population in control, siAND1- and siRAD21-treated S-phase cells. Note the reduced enrichment of the chromatin-bound DH population in siAND1-treated cells (p = 0.08) and siRAD21-treated cells (p ≤ 0.01) relative to S phase controls. **e.** Box plot showing enrichment of chromatin-bound SH population in control, siAND1- and siRAD2-treated S-phase cells. Note no obvious change in SH population in siAND1 (p = 0.95) and notable change in SH population for siRAD21 (p = 0.06) relative to S-phase controls. **f.** Representative MINFLUX images of siRAD21-treated cells respectively. MINFLUX confocal scan of S phase cell. Top left: MCM2-Halo (red); Top right: 10-minute pulse EdU positive cell (cyan, scale bar 5 µm); inset: chosen ROI for MINFLUX; Bottom left: MINFLUX localizations overlay with confocal ROI; Bottom right: MINFLUX localizations (yellow, scale bar 200 nm). (inset in top row: ROI for MINFLUX imaging). **g.** Histograms of nearest-neighbor distances (nm) between MCM2 molecules for compiled S-phase data from siRAD21-treated cells (teal) with the simulated dimeric distribution (orange) and untreated S-phase population (black line). **h.** Box plot showing enrichment of the DH population in G1 and S-phase in the presence and absence of RAD21. Note that a significant change is observed only during S-phase. Statistical analyses are included in Supplementary table 4.

To quantify the inter-replisome distance, we subtracted the simulated dimer distribution from the observed NN distributions to fit the residual signal with a Gaussian model, isolating the coupled-replisome population. In unperturbed S-phase cells, this yielded an inter-replisome distance of 39 ± 5 nm (Fig. 5c, Extended Fig. 8e). Strikingly, AND1 depletion expanded this distance to 53 ± 5 nm (Fig. 5c), demonstrating that AND1 directly constrains sister replisomes within a defined spatial proximity of ∼40 nm.

To test whether higher-order chromatin architecture further contributes to replisome organization, we depleted RAD21, a core structural subunit of the cohesin complex (Fig. 5f-g, Extended Fig. 7b). Cohesin is enriched at replication origins and interacts with MCM proteins^32^, and its loss has been linked to altered origin firing dynamics, though effects vary by cell type^32^. RAD21 depletion in MCM2-Halo U2OS cells altered neither total MCM levels nor global cell cycle distribution (Fig. 5f, Extended Fig. 7b-c). Crucially, MCM DH organization in G1 was indistinguishable from controls (Fig. 5g-h, Extended Fig. 8b-c), suggesting that MCM loading is independent of cohesin. During S-phase, however, RAD21 depletion caused a significant reduction in the DH population (Fig. 5d) and produced a broad, multimodal NN distribution at distances >60 nm (Fig. 5g, Extended Fig. 8d). This multimodal pattern contrasts sharply with the defined Gaussian shift observed upon AND1 depletion. Rather than a uniform increase in inter-replisome distance, cohesin loss produced a heterogeneous population of MCM separations, consistent with a breakdown of organized spatial confinement rather than specific uncoupling of the sister CMG pair itself.

Together, these findings reveal two mechanistically distinct contributions to replisome coupling: AND1-mediated physical tethering of sister CMGs at a defined ∼40 nm proximity, and cohesin-dependent chromatin architecture that organizes replisomes within higher-order nuclear domains during S-phase (Fig. 6).

**Figure 6.**
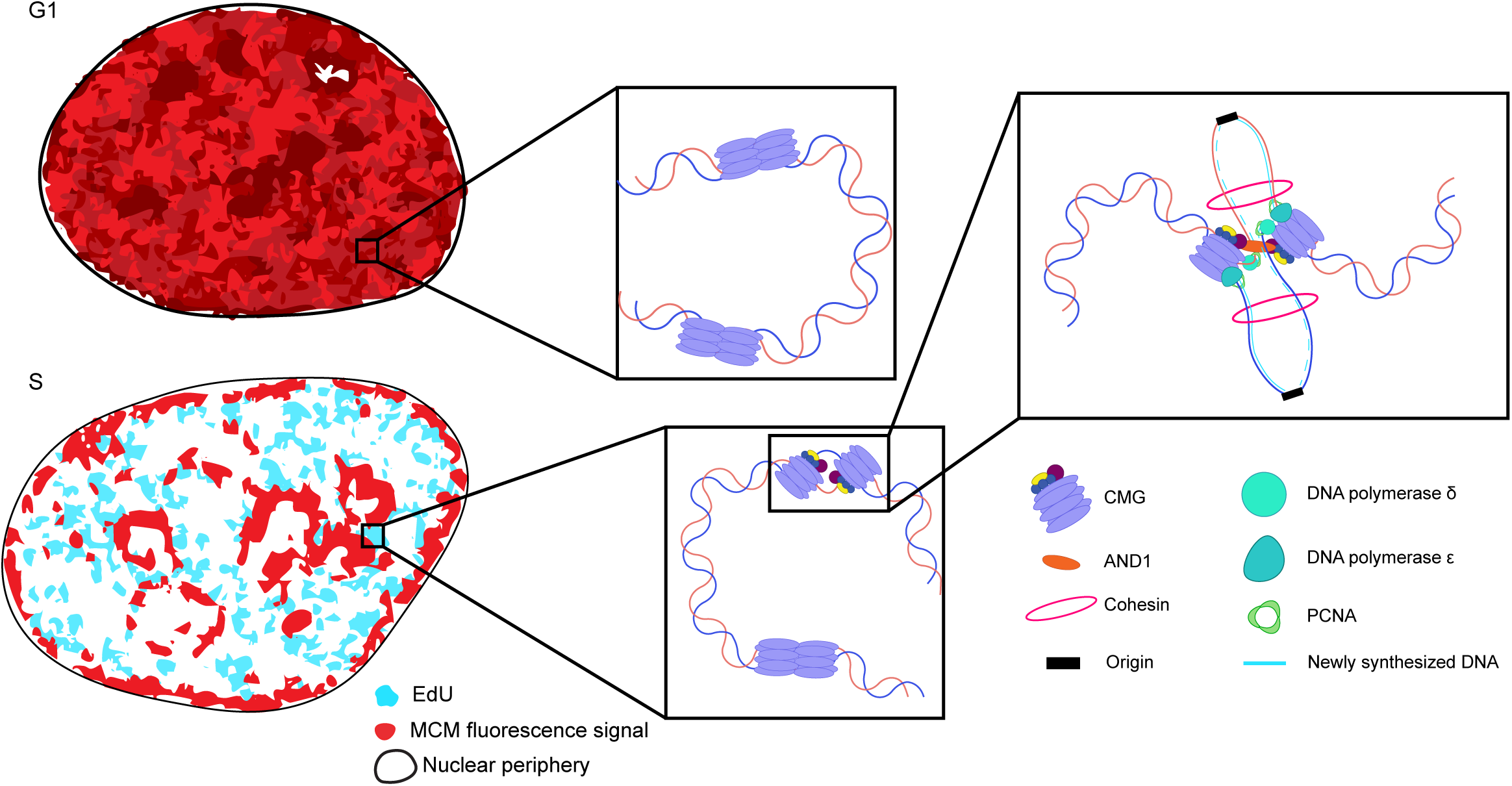
Schematic representation of DH and SH of MCMs during G1 and S-phase. Note the coupling of replisomes by AND1.

## DISCUSSION

Although biochemical and cryo-EM studies using purified proteins established the existence of MCM DHs and proposed how they transition into single hexamers (SHs) during replication, direct spatial evidence within intact mammalian cells has been lacking. Here, we provide the MINFLUX-based visualization of the DNA replication machinery in the native cellular context, delivering direct spatial observation of the DH-to-SH transition during S-phase (Fig. 6).

A major advance toward understanding DNA replication in its native context came from the demonstration that mammalian replication foci can be resolved as individual replicons rather than clusters of replicons^51^. More recently, super-resolution imaging of newly synthesized DNA has revealed the spatial organization and dynamics of active replication nanostructures^52^. Our study extends this structural mapping directly to the protein machinery itself. By visualizing the in situ architecture of the replisome, we link the nanoscale organization of DNA synthesis sites to the physical arrangement of the replication machinery.

By resolving MCM DH and SH states, MINFLUX enables us to distinguish active from dormant MCM complexes and quantitatively define origin activation in situ. We estimate that, on average during an unperturbed S phase, only approximately 8% of chromatin-bound MCMs are in the active SH forms (Supplementary Note 1), providing direct structural confirmation that the vast majority of loaded MCMs serve as an inactive, dormant reserve. Despite mobilization of this dormant reserve under replicative stress, our quantification suggests that the number of active replisomes remains limited. For instance, despite a marked increase in CDC45 chromatin engagement following ATR inhibition, indicative of enhanced CMG assembly and origin firing, the corresponding increase in markers of active DNA synthesis is comparatively modest at approximately 1.4-fold^50^. Scaling our measured ∼8% active replication forks by this 1.4-fold increase in DNA synthesis yields an estimated ∼11.2% active forks upon ATR inhibition, which closely matches the ∼11% value calculated independently from our imaging data (Supplementary Note 1). The agreement between these estimates argues against a simple runaway increase in origin firing alone. The increased helicase assembly does not translate proportionally into productive DNA synthesis, supporting a model where productive fork establishment is constrained by the finite availability of replication factors required downstream of helicase activation. This is consistent with emerging evidence that the S-phase checkpoint restrains origin activation to prevent replisome component depletion^53–55^.

We note that these quantifications likely represent conservative lower-bound estimates because our measurements are based on two-dimensional projections; the three-dimensional orientation of MCM complexes can influence the observed spatial distributions. Consequently, larger nearest-neighbor distances measured in 2D may occasionally reflect chance proximities to MCMs from a different origin rather than true uncoupling, particularly if a DH partner lies in an axial orientation. Despite these technical constraints, MINFLUX clearly captures the active splaying of MCM DHs into SHs.

Strikingly, even as the two hexamers physically separate, the resulting sister replisomes do not freely diffuse apart, instead, they remain tethered at a constant inter-replisome distance of ∼40 nm throughout S-phase. This persistent spacing indicates that bidirectional fork progression is spatially constrained, keeping sister replisomes functionally anchored with a shared replication factor or higher-order chromatin domain-regardless of replication stage. Our MINFLUX measurements find independent support from recent genome-scale studies of replication fork organization. Liu et al. used replication-associated Hi-C (Repli-HiC) to identify fountain-like chromatin contact structures centered at replication initiation zones, with a median length of ∼160 kb, interpreted as the genomic footprint of coupled sister forks^22^. Polymer simulations suggest that two loci separated by 160 kb on chromatin would be ∼500 nm apart in three dimensions if the forks were uncoupled, which is far beyond the effective contact range of Hi-C ligation. The existence of the fountain therefore implies that sister forks are held in close physical proximity throughout elongation, consistent with the ∼40 nm inter-replisome distances we measure directly by MINFLUX.

To dissect the molecular mechanisms underpinning this replisome coupling, we perturbed two distinct spatial organizers: AND1 and cohesin. Depletion of AND1 (the human Ctf4 ortholog), a trimeric scaffold proposed to link multiple CMGs within replication factories, significantly expanded the inter-replisome distance. This provides direct structural support for a model where AND1 physically tethers sister replisomes at a local, molecular level. Conversely, to test the role of chromatin-based coupling, we depleted RAD21, a core kleisin subunit of the cohesin complex. As replication forks advance, cohesin-mediated spatial confinement appears critical for keeping sister forks within the same 3D territory. Our observation that RAD21 loss drastically reduces the DH population and yields a highly heterogeneous distribution of SH separations strongly implicates higher-order chromatin architecture in replisome coupling. This aligns with recent models proposing that cohesin-mediated loop anchors confine human replication origins^31^, potentially pushing licensed MCMs in G1 to localize and activate specific origins at topological boundaries^30^.

Two orthogonal approaches, our nanometer-scale single-molecule localization shown here and genome-wide chromatin contact mapping from Liu et al., converge on the same conclusion: sister replisomes are physically coupled throughout S-phase, with AND1 serving as a key molecular mediator of this spatial constraint.

More broadly, the spatial organization of bidirectional DNA replication has long been framed as a strict dichotomy. On the one hand, biochemical, structural, and single-molecule biophysical assays indicate that sister CMG helicases must physically uncouple upon origin firing to function as autonomous, independent tracking motors capable of bypassing structural lesions. On the other hand, genomic contact mapping and conventional live-cell imaging strongly support a macroscopic replication factory model, wherein sister forks remain physically coupled. Our MINFLUX measurements resolve this paradox by revealing that sister replisomes separate but do not freely diffuse apart, maintaining a characteristic spatial separation of ∼40 nm throughout S-phase. This ∼40 nm gap provides the necessary steric clearance for independent motor function and topological bypass, fully satisfying the mechanical constraints of the uncoupled model. Simultaneously, because this distance is well below the diffraction limit of standard light microscopy and highly confined within genomic interaction maps, it perfectly explains why macroscopic assays observe a rigidly coupled factory. Ultimately, our data demonstrates that sister replisomes are neither fused nor independent; rather, they are autonomously functioning machines maintained within a tightly constrained ∼40 nm spatial hub by the combined forces of AND1 tethering and cohesin-dependent chromatin architecture. By directly visualizing this molecular architecture in situ, this approach resolves a long-standing structural paradox and opens powerful new avenues for interrogating DNA replication dynamics within their native cellular context.

AI tools were used to improve grammar and conciseness.

## Acknowledgements

We thank members of the Prasanth laboratory for discussions and suggestions. We thank Drs. Bruce Stillman, Jessica Matthias, Chris Zwilling for providing reagents, suggestions and for technical help. We thank the IGB Microscopy core, Glenn Fried, Kingsley A. Boateng, and Reza Rajabi Toustani for providing advice, reagents and technical help. We thank Abberior staff Jessica Matthias for providing advice, sample preparation tips and sample reagents. This work was supported by NSF-QCB-STC (NSF DBI 2243257) to OJZ, TH, KVP, SGP, ARPA-H (AY1AX000030) to KVP and NIH R01GM132458 to KVP; R35GM122569 to TH, and NSF (2225464) and NIH (1R35GM152450) awards to SGP. JZ was supported by a Feodor Lynen Research Fellowship of the Alexander von Humboldt Foundation. HPS was supported by the Czech Science Foundation Junior Star (grant 22-20303M), the EU Horizon 2022 Widera Talent programme (ERA grant agreement 101090292) and an EMBO installation grant (IG-5689-2024). TH is an investigator with the Howard Hughes Medical Institute.

## Author Contributions

OJZ designed and performed most experiments; JZ analyzed most experiments. HPS provided the MCM2/4-Halo lines. SGP, KVP and TH supervised the project. OJZ, JZ, TH and SGP wrote the manuscript. All authors read the manuscript and provided input.

## Ethics declarations

## Abbreviations

MCM: Mini-Chromosome Maintenance
MINFLUX: MINimal photon FLUX
PCNA: Proliferating Cell Nuclear Antigen
EdU: 5-Ethynyl-2’-deoxyuridine
SH: single hexamer
DH: double hexamer
HU: hydroxyurea

## Extended Figure Legends

**Extended Figure 1.**
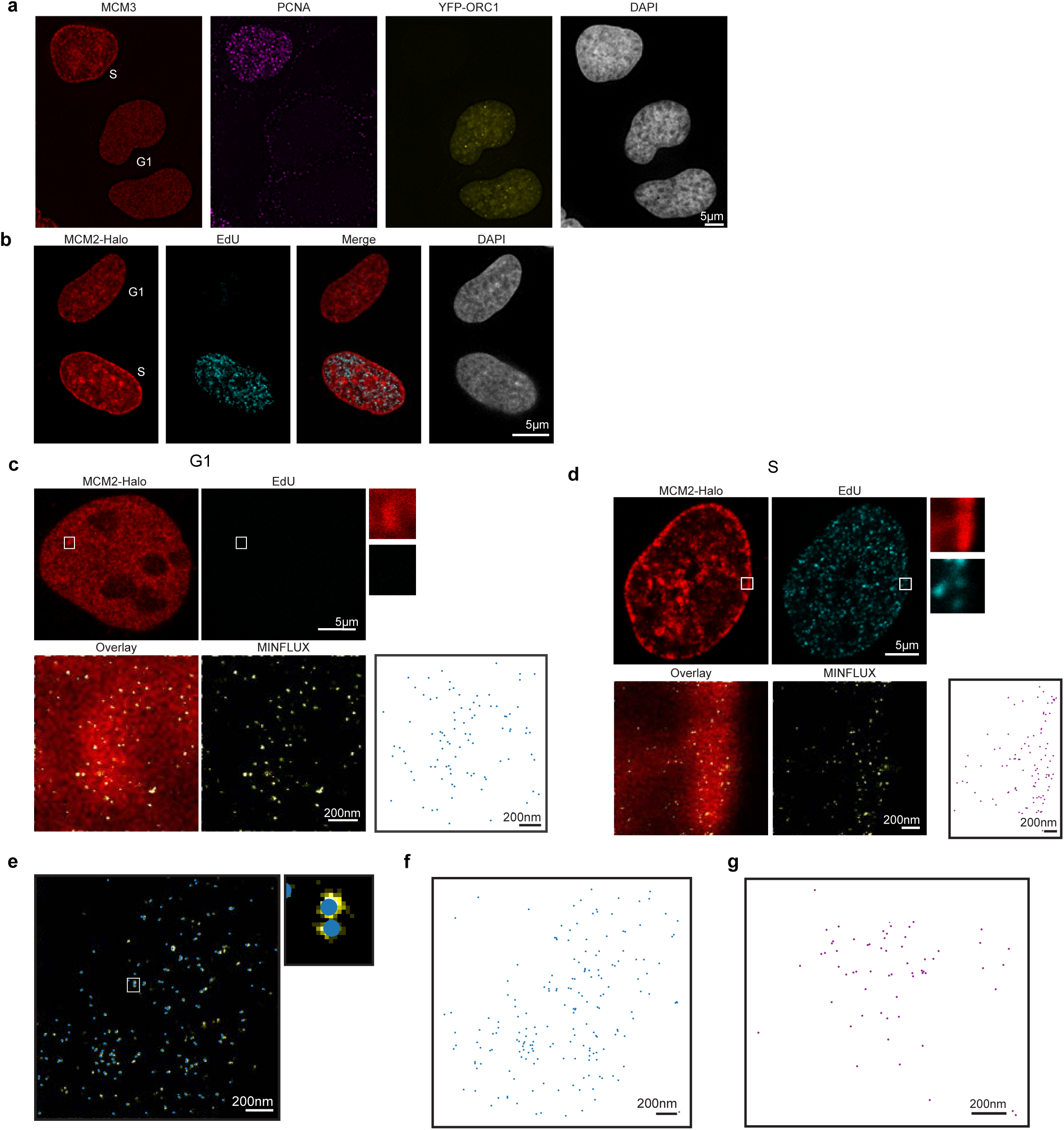
Visualizing MCM using MINFLUX. **a.** Immunostaining of MCM2 (red) with PCNA (magenta) in YFP-ORC1 (yellow) stable U2OS cells. Top cell in S-phase (PCNA-positive, ORC1-negative), bottom two cells are in G1 phase (YFP-ORC1-positive and PCNA-negative). **b**. MCM2-Halo (red) cells with EdU labeling (cyan, 10-minute pulse). Top cell is in G1 (no EdU signal), bottom cell is in S phase (EdU signal present). **c.** Additional representative G1 phase cell. **d.** Additional representative S-phase cell. **e.** MCM2-Halo MINFLUX localizations from Figure 1f & h overlayed with dot plot of identified molecules after analysis. Inset of double hexamer from Figure 1f & h. **f-g.** Full plot of MCM localizations from Figure 1h-i, expanded to include all identified MCM (not confined due to Imspector software restrictions).

**Extended Figure 2.**
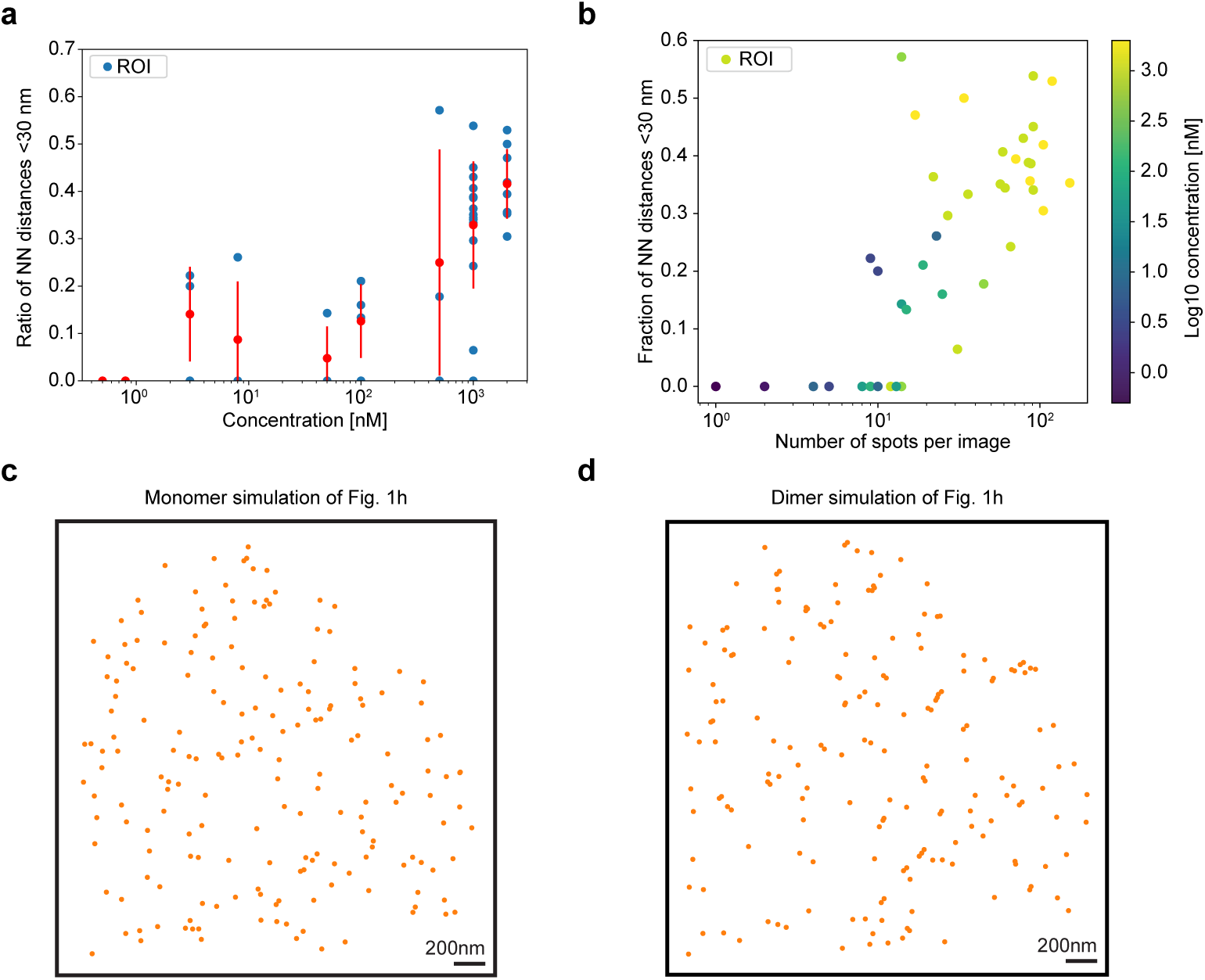
Quantification and simulation analysis of MCM localization. **a.** Ratio of the nearest neighbor of spots within each ROI identified increasing as an increase in FLUX 640 Halo ligand concentration. **b.** Fraction of nearest neighbor distances <30 nm within each ROI increasing as number of spots identified increases with higher log10 concentration of FLUX 640 Halo ligand (blue to yellow). Note the intermingling of light green and yellow spots, indicating saturation of ligand. **c-d.** Monomer and dimer simulation re-distribution of identified molecules in MINFLUX image from Fig. 1h respectively.

**Extended Figure 3.**
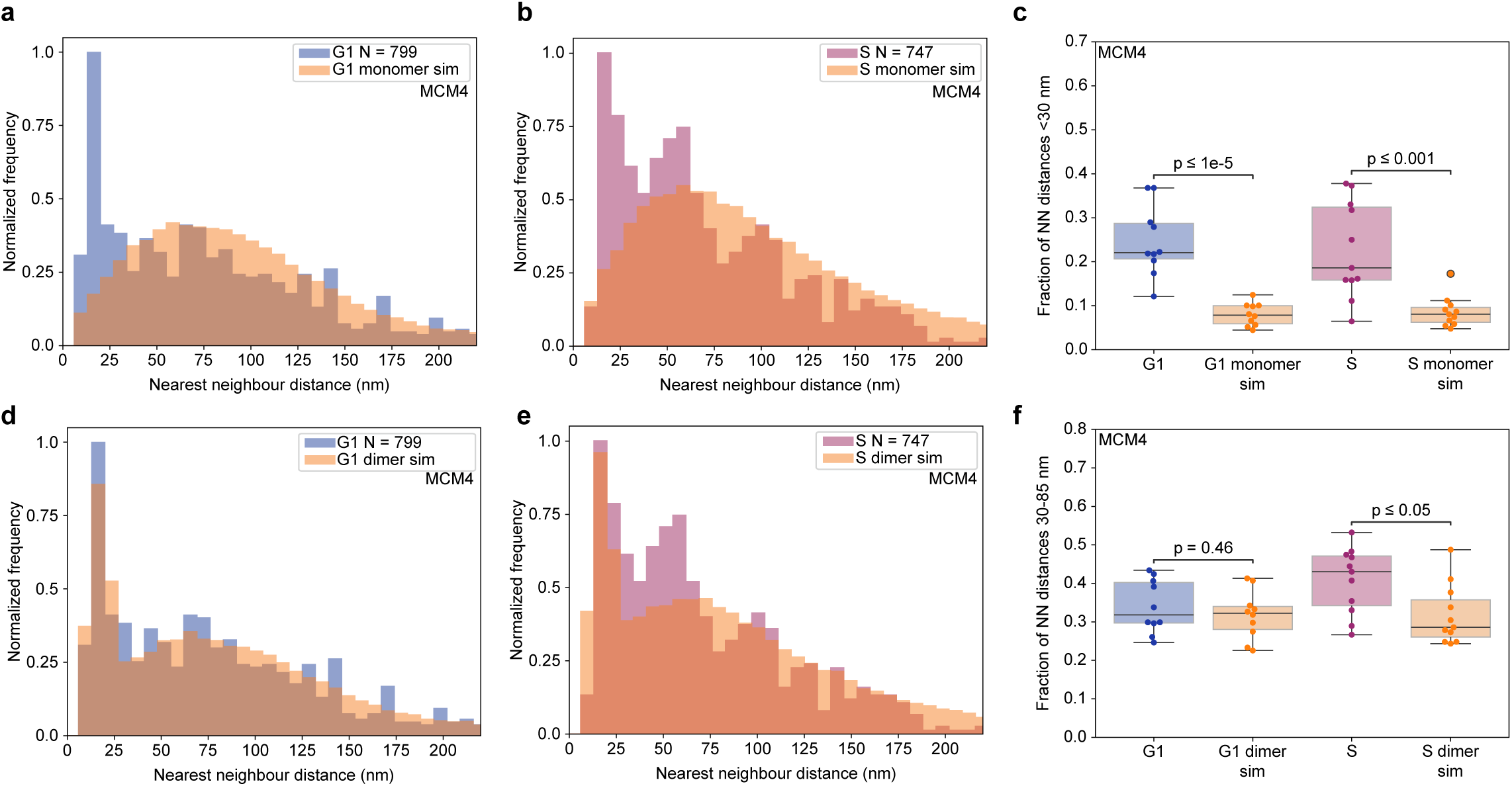
Visualizing MCM4 localization. **a-b**. Histograms of nearest neighbor distance (nm) between MCM4 molecules for compiled G1 and S-phase data, respectively compared to monomer simulation distributions. **c**. Box plot depicting chromatin-bound DH of MCM4 population in G1 (p ≤ 0.001) and S (p ≤ 1e-5) compared to monomer simulation distribution **d-e.** Histograms of nearest-neighbor distances (nm) between MCM4 molecules for compiled G1 and S-phase data, respectively, compared with dimer simulated distributions. **f**. Box plot depicting chromatin-bound SH of MCM4 population for G1 (p=0.60, non-significant), and in S (p ≤ 0.001) compared to dimer simulation distribution.

**Extended Figure 4.**
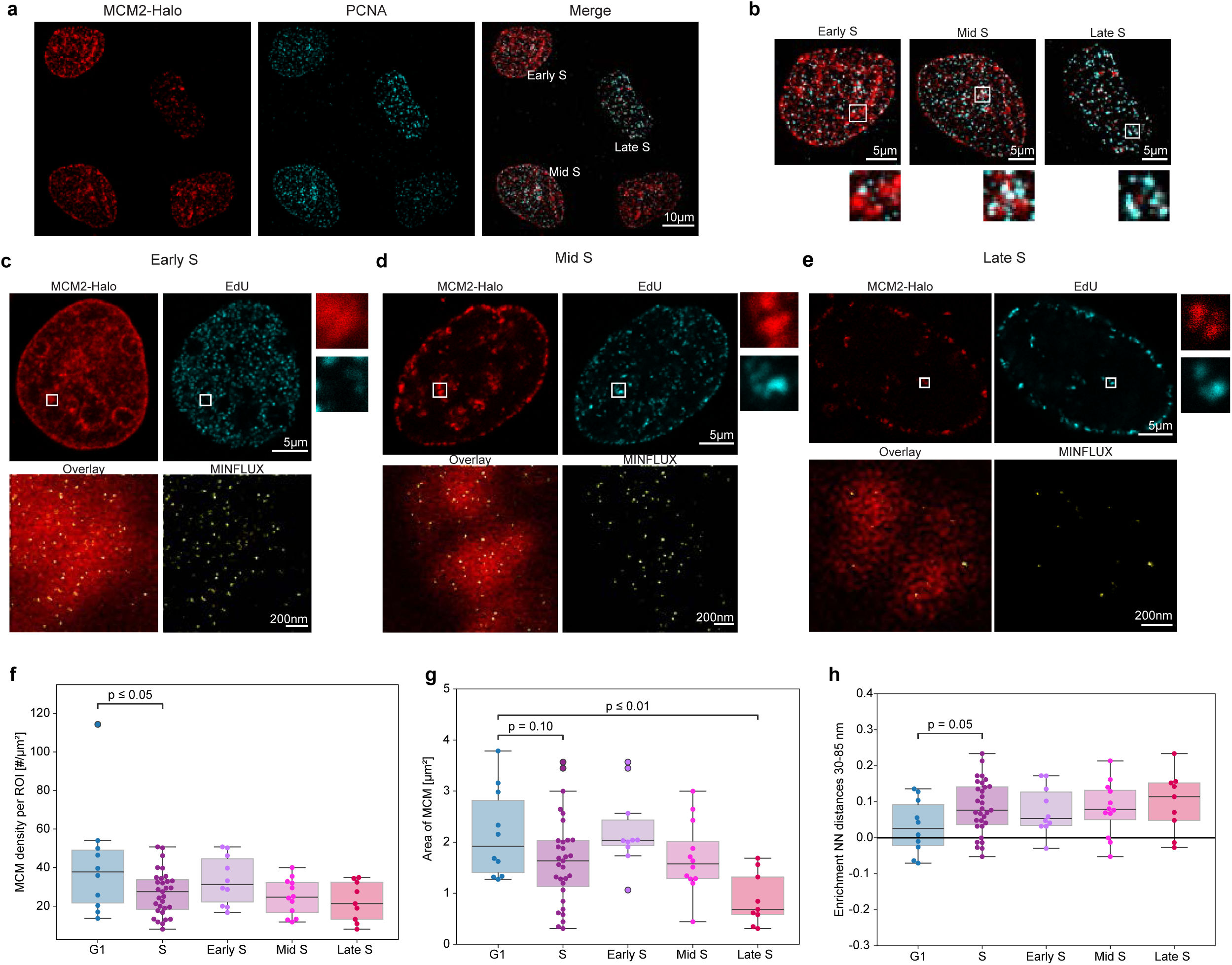
Spatial organization of MCM localization across S-phase. **a-b.** Immunostaining of PCNA (cyan) with MCM2-Halo (red) showing distinct spatio-temporal patterning for both MCM and PCNA. The insets are showing a change in patterns from early, mid and late S-phase. Note the lack of overlap between MCM and PCNA by conventional microscopy. **c-e.** Representative MINFLUX confocal scans of early, mid, and late S phase cell. Top left: MCM2-Halo (red); Top right: 10-minute pulse EdU positive cell (cyan, scale bar 5µm); inset: chosen ROI for MINFLUX; Bottom left: MINFLUX localizations overlay with confocal ROI; Bottom right: MINFLUX localizations (yellow). **f.** Box plot showing the number (#) of chromatin-bound MCM per micron squared. A significant decreasing from G1 to S (p ≤ 0.05) followed by the progression of decreasing density from early, to mid to late S-phase is observed. **g.** Box plot showing area of chromatin-bound MCM. Note a significant decrease from G1 to S (p ≤ 0.05) followed by the progression of decreasing area from early to mid to late S-phase. **h.** Box plot depicting SH in G1. Note the significant change of enrichment of chromatin-bound SH population in G1 to S (p=0.05) followed by the progression of increased distances from early, to mid to late S-phase.

**Extended Figure 5.**
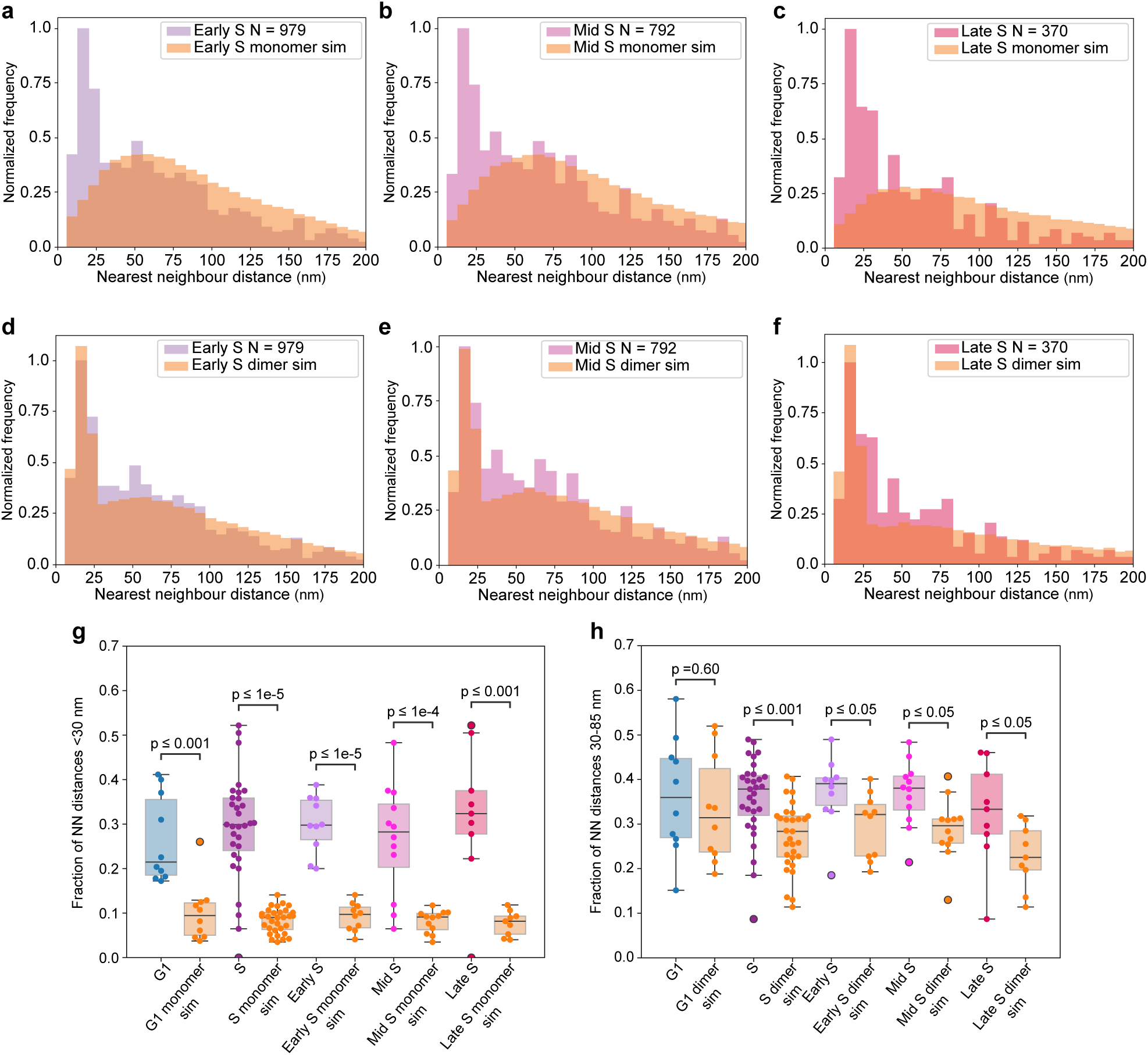
Progression of MCM localization across S-phase. **a-f.** Histograms of compiled early, mid, and late S-phase data compared to monomer and dimer simulation distribution, respectively. **g.** Box plot showing chromatin bound DH population in G1 and throughout S phase. Note the significant enrichment of the chromatin-bound DH population in G1 (p ≤ 0.001) S phase (p ≤ 1e-5), early S-phase (p ≤ 1e-5), mid S-phase (p ≤ 1e-4), and late S-phase (p ≤ 0.001) compared with the monomer simulation distribution, whereas no significant enrichment is observed in G1 phase (p = 0.60). **h.** Box plot showing chromatin bound SH population in G1 and throughout S-phase. Note the significant enrichment of the chromatin-bound SH population in S phase (p ≤ 0.001), early S-phase (p ≤ 0.05), mid S-phase (p ≤ 0.05), and late S-phase (p ≤ 0.05) compared with the dimer simulation distribution, whereas no significant enrichment is observed in G1 phase (p = 0.60).

**Extended Figure 6.**
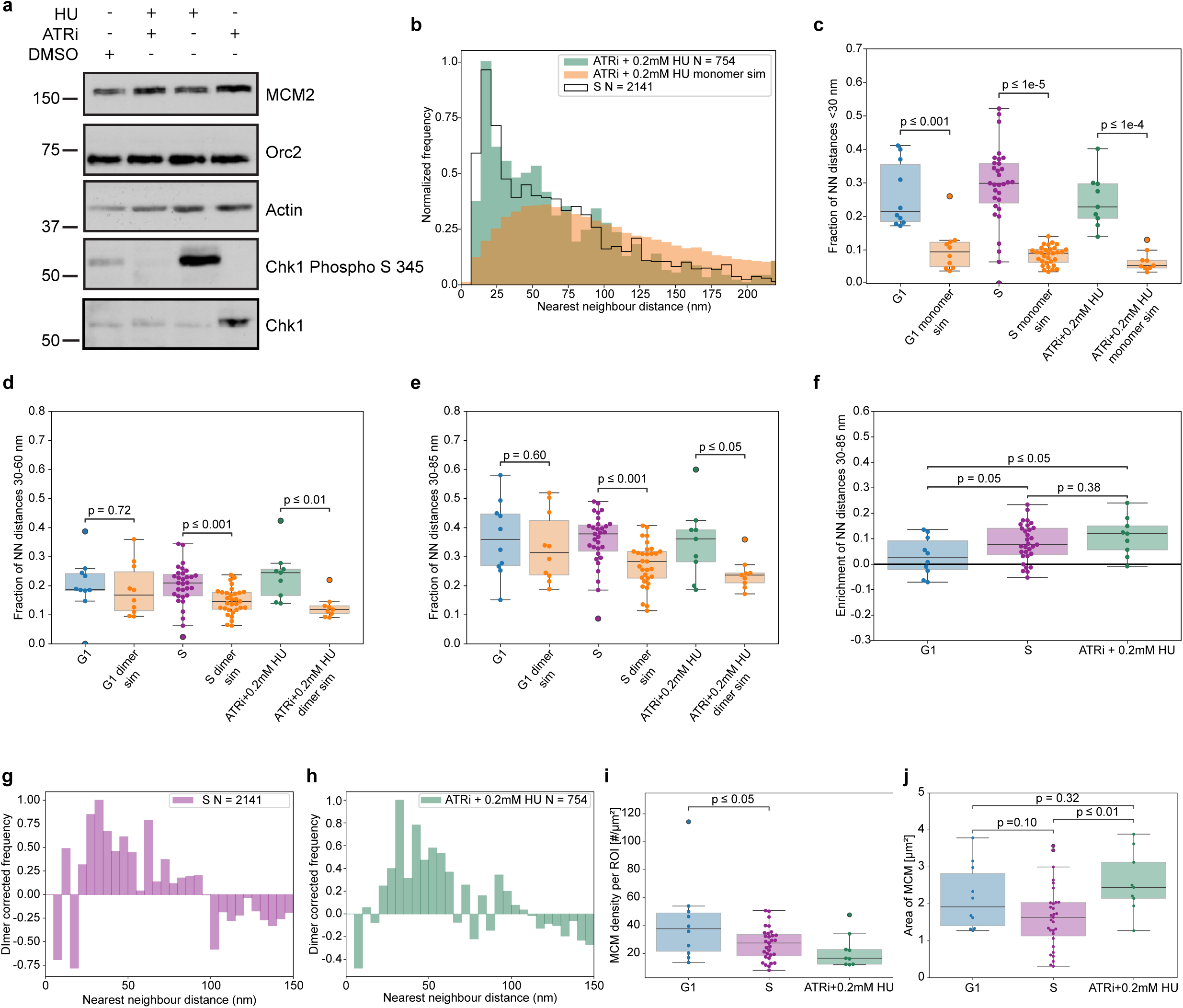
Replication stress and ATR inhibition alter the spatial organization of MCM complexes during S phase. **a.** Western blot with various antibodies (MCM2, Orc2, Actin, Chk1 phospo-345, Chk1) in MCM2-Halo cells treated with ATRi+0.2 mM HU. Note an increase of Chk1 Phosphorylation at S345 with 0.2 mM HU alone, confirming replication stress. ATR inhibition in presence of HU showed no change in Chk1 Phosphorylation at S345 confirming inhibition of ATR. **b.** Histogram of nearest neighbor distance (nm) between MCM2 molecules for compiled ATRi+0.2 mM HU treated S phase data (green) compared to monomeric simulation distribution (orange) and untreated S phase population (black line). **c.** Box plot depicting DH in G1, S and ATRi+0.2 mM HU. Note the significance of chromatin-bound DH population in G1 (p ≤ 0.001), S (p ≤ 1e-5), and ATRi+0.2 mM HU (p ≤ 1e-4). **d.** Box plot of SH population in G1, S and ATRi+0.2 mM HU (30-60 nm). Note the significance of chromatin bound SH population in S (p ≤ 0.001), and ATRi+0.2 mM HU (p ≤ 0.01) and non-significance of chromatin bound SH population for G1 (p=0.72) **e.** Box plot of SH population in G1, S and ATRi+0.2 mM HU (30-85 nm). Note the significance of chromatin bound SH population in S (p ≤ 0.001), and ATRi+0.2 mM HU (p ≤ 0.05) and non-significance of chromatin bound SH population for G1 (p=0.60) **f.** Box plot showing chromatin-bound SH population (30-85 nm) from G1 to S and ATRi+0.2 mM HU treated. Note the significant enrichment of SH population from G1 to S and to ATRi+0.2 mM HU and a notable increase in SH population in ATRi+0.2 mM HU treated compared to S-phase cells. **g-h.** Dimer corrected frequency histograms of S-phase and ATRi+0.2 mM HU. **i.** Box plot showing the number (#) of chromatin-bound MCM per micron squared. Note the decrease in G1, S (p ≤ 0.05), and ATRi+0.2 mM HU. **j.** Box plot showing area of chromatin-bound MCM. Note the significant decrease from G1 to S (p ≤ 0.05) and ATRi+0.2 mM HU (p = 0.94, G1 to ATRi+0.2 mM HU and p ≤ 0.05 S to ATRi+0.2 mM HU).

**Extended Figure 7.**
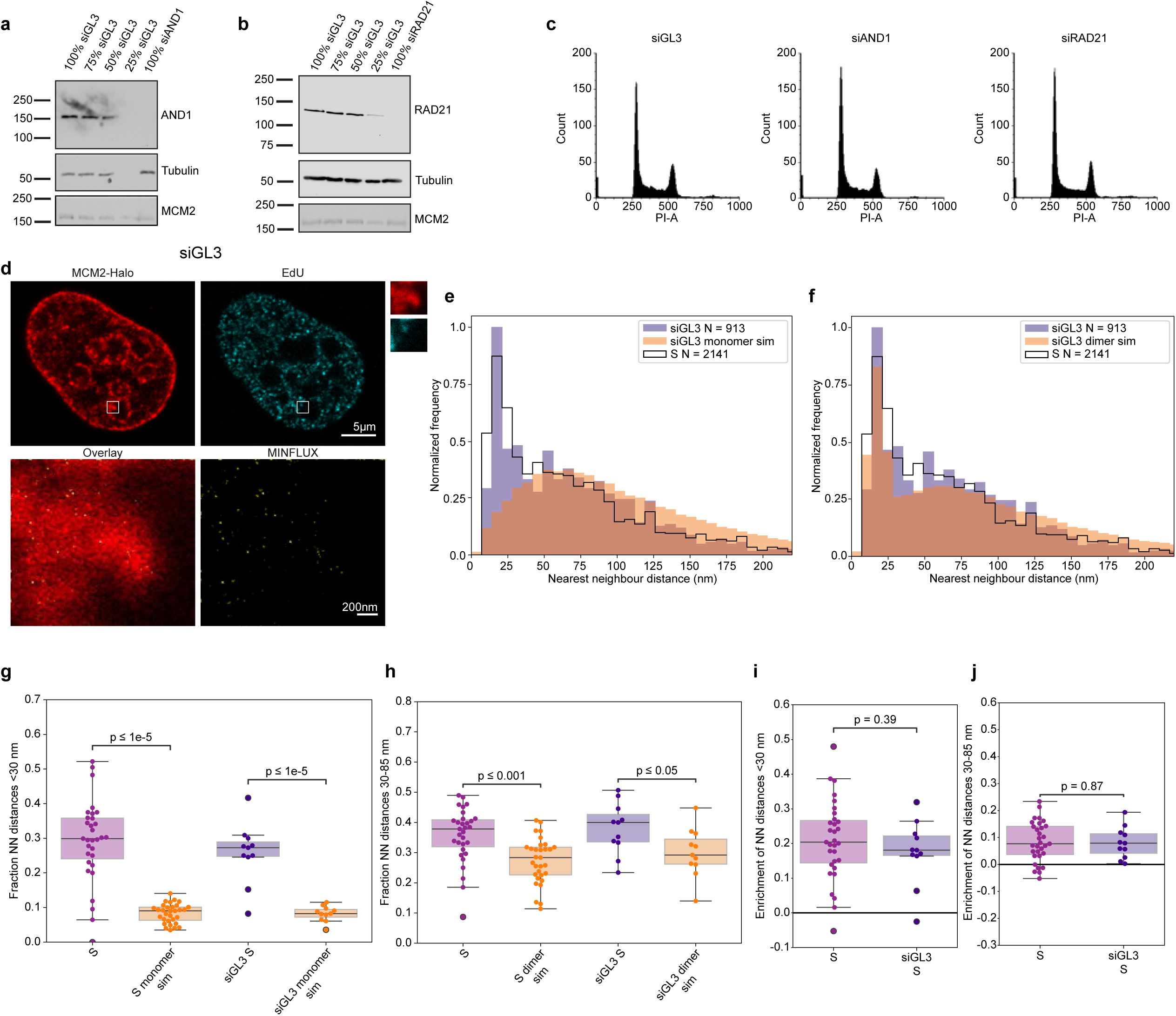
Nanoscale organization of chromatin-bound MCM2 in control and siGL3-treated cells. **a.** Western blot in control (siGL3), siAND1-treated MCM2-Halo cells using AND1, Tubulin and MCM2 antibodies. **b.** Western blot in control and siRAD21-treated cells using RAD21, Tubulin and MCM2 antibodies. **c.** PI flow of siGL3-, siAND1-, siRAD21-treated MCM2-Halo cells. **d.** Representative MINFLUX confocal scan of siGL3 treated S phase cell Top left: MCM2-Halo (red); Top right: 10-minute pulse EdU positive cell (cyan, scale bar 5 µm); inset: chosen ROI for MINFLUX; Bottom left: MINFLUX localizations overlay with confocal ROI; Bottom right: MINFLUX localizations (yellow, scale bar 200 nm) **e-f.** Histogram of nearest neighbor distance (nm) between MCM2 molecules for compiled siGL3-treated S phase data (purple) compared to monomer and dimer simulation distribution (orange), respectively, and untreated S phase population (black line). **g.** Box plot showing DH population during S-phase in control and siGL3-treated cells. Note the significance of chromatin-bound DH population in siGL3 (p ≤ 1e-5) compared to monomeric simulation distribution. **h.** Box plot showing SH in S-phase control and siGL3. Note the significance of chromatin-bound SH population in siGL3 (p ≤ 0.05) compared to dimeric simulation distribution. **i.** Box plot showing chromatin-bound DH population from S-phase to siGL3 (p = 0.39). **j**. Box plot showing enrichment of chromatin-bound SH population in S-phase compared to siGL3 (p=0.87).

**Extended Figure 8.**
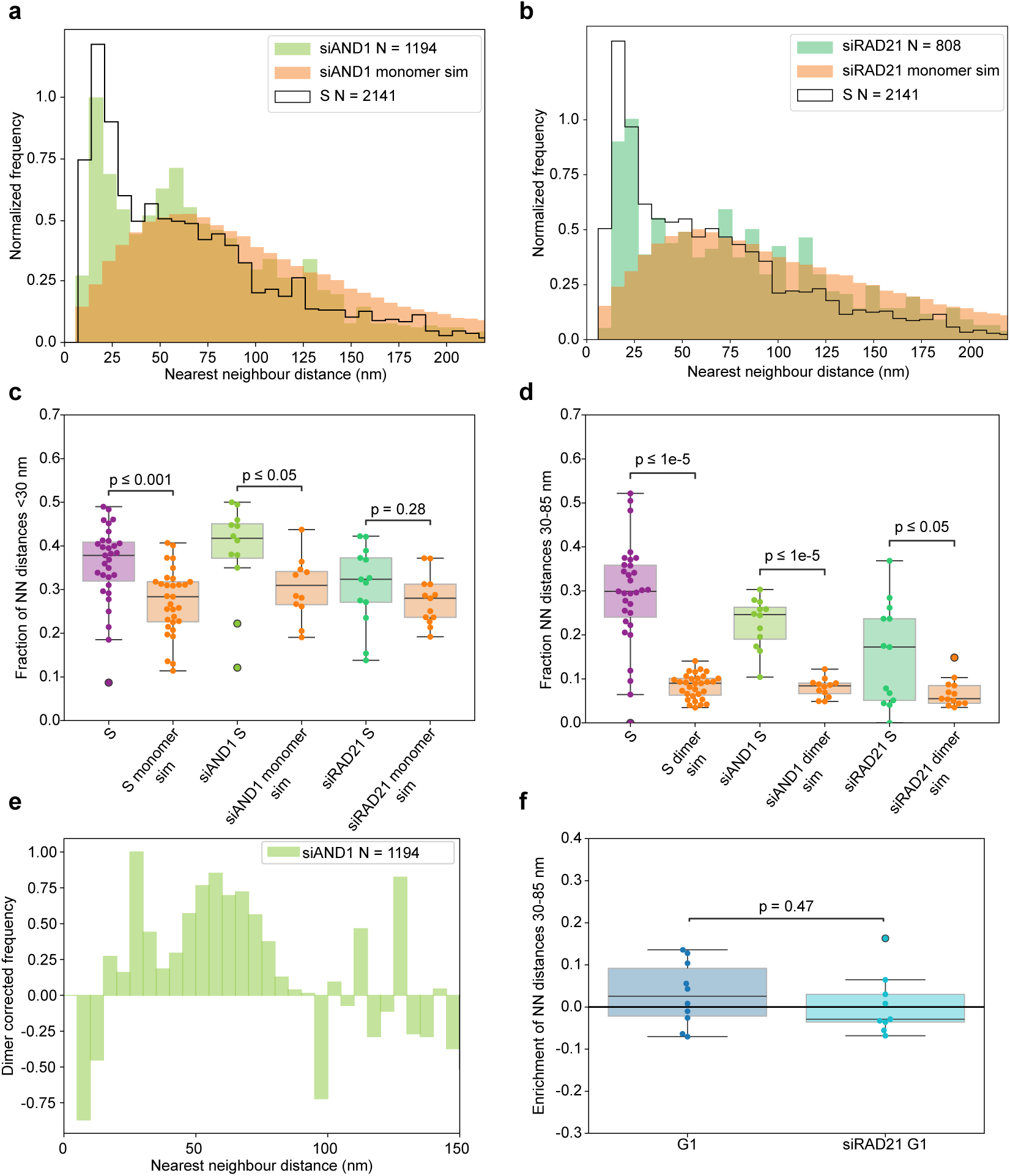
Perturbation of AND1 and RAD21 alters the nanoscale architecture of chromatin-bound MCM2 complexes. **a-b.** Histogram of nearest neighbor distance (nm) between MCM2 molecules for compiled siAND1- and siRAD21-treated S-phase data (green, teal respectively) compared to monomeric simulation distribution (orange) and untreated S-phase population (black line). **c.** Box plot depicting DH in S, siAND1-, and siRAD21-treated S-phase data. Note the significance of chromatin-bound DH population in siAND1, (p ≤ 1e-5), and siRAD21 (p ≤ 0.05). **d.** Box plot of SH population in S, siAND1-, and siRAD21-treated S-phase data. Note the significance of chromatin bound SH population in siAND1, (p ≤ 0.05), and non-significance of siRAD21 (p = 0.28). **e.** Dimer corrected frequency histogram of siAND1. **f.** Box plot showing enrichment of chromatin-bound SH population in G1 compared to siRAD21-treated cells in G1 (p =0.47).

---

**Supplementary Note 1**

To estimate the active SH form, we calculated the enrichment of the MINFLUX nearest-neighbor (NN) distribution over the simulated monomer and dimer distributions per ROI. In S phase, an enrichment within the 30–85 nm window was observed. Identifying this population as the SH form is supported by the fact that all measurements of G1-phase cells showed no significant enrichment (MCM2: 3 ± 2%; MCM2 with siRAD21 depletion: 0 ± 2%; MCM4: 2 ± 2%) (Fig. 2h, 2j, Extended Fig. 8f). Additionally, after activating dormant origins (ATRi+0.2 mM HU) we observed a more distinct peak in the NN distance and an increase in enrichment within the 30–85 nm range (Fig. 4c). Lastly, MINFLUX measurements of MCM4 also exhibited a distinct peak in this same region (Fig. 3f). Therefore, we used the enriched NN fraction as an estimate for the SH fraction. The mean enrichment across MCM foci during S phase was 8.4 ± 1.3% for MCM2, 9 ± 3% for MCM4, and 10.8 ± 1.7% for MCM2 with ATRi+0.2 mM HU. %) (Fig. 2h, 2j and Extended Fig. 6f). No corrections were performed for 3D orientation or labeling fraction. Note that the estimates are only snapshots of active MCM at a given time, thus will bias it towards it being a lower estimate. All values are presented as mean ± SEM.

