## Supplementary material for "Direct visualization of MCM helicase activation and replisome coupling in situ": Methods

**Materials and Methods:**

**Cell Culture:**

MCM2/4-Halo U2OS cells were grown in Dulbecco's modified Eagles medium (DMEM) supplemented with 5% fetal bovine serum (FBS) and 1% pen strep (P/S). YFP-ORC1 stable U2OS cells were grown in Dulbecco's modified Eagles medium (DMEM) supplemented with 5% FBS, 1% P/S and 800 ug/ml G418.

Drug treatments:

Cells were incubated with 0.2 mM HU and 10 µM ATRi (NU6027) for 4 hours at 37°C. Cells were then stained using MINFLUX sample prep or lysate collection for western blot analysis in experiments indicated.

Knockdowns:

Cells were grown to 40-50% confluency. Media was changed to OptiMEM (ThermoFisher CAT# 11058-021), 10 nM siRNA (siAND1 (L2-019780-01-0005), siRAD21 (L2-006832-01-0010)) SMARTpool mixed with Lipofectamine RNAiMAX (ThermoFisher CAT# 13778150) incubated for 5 hours, then complete media was added for 24 hours. Cells were then stained using MINFLUX sample prep, lysate collection for western blot analysis, or PI flow in experiments indicated.

**MINFLUX sample preparation:**

Cells were plated onto 1.5mm coverslips and grown to 50-75% confluency. Cells were incubated with 125 µM EdU for 10 minutes. Media with EdU was washed off once with CSK buffer and cells were pre-extracted with 0.5% Triton X-100 in CSK buffer for 5 minutes on ice. Cells were then fixed in 2% PFA for 30 minutes, followed by a 5-minute quench with 50 mM NH4Cl. Cells were washed twice with 1X PBS for 5 minutes. Click-It solution made in 1X PBS containing final concentration 100 mM Na-ascorbate, 10 mM $CuSO_4$, and 1 µM 488-azide was incubated on the coverslip for 1 hour in a covered humidified chamber. Cells were washed twice with 1X PBS for 5 minutes. Cells were then blocked with Image-iT Signal Enhancer (Thermo CAT# I36933) for 30 minutes in a covered humidified chamber. Cells were rinsed with 1X PBS then immediately incubated with Halo ligand in 0.5-1% BSA in 1X PBS for 1 hour in a covered humidified chamber. (ligand concentrations ranging from 800 pM to 2 µM, 1 µM and 2 µM were used for data analysis). Cells were rinsed 3 times with 1X PBS then washed 3 times with 1X PBS for 5 minutes each. Coverslips were incubated with 100 µl gold beads (BBI CAT# EM.GC150/7) for 15 minutes at room temp, followed by several washes with 1X PBS to remove non-bound beads. Coverslips were mounted on slides (GSC CAT# 4-13057-DZ-12) with GLOX imaging buffer (50 mM TRIS/HCl, 10 mM NaCl, 10% glucose, 0.4 mg/ml Glucose Oxidase (Sigma G2133-50KU) in Tris 63 µg/ml catalase (Sigma C1354-1G) in $H_2O$, 10-25 mM MEA (M6500-25G), pH 8.0).

**Imaging sequence:**

For the MINFLUX imaging sequence a hexagonal 2D MINFLUX pattern and a stickiness parameter of 2 were chosen. .

**Immunofluorescence:**

Cells were plated onto 1.5mm coverslips and grown to 50-75% confluency. If Halo ligand was incubated pre-fix: 1 nM Halo Ligand JFX 585 (Promega REF# HT1040) or Halo Ligand JFX 650 (Promega REF# HT1070) incubation 12-16 hours in cell culture media. If EdU was used, cells were incubated with 125 μM EdU for 10 minutes. Media with EdU was washed off once with CSK buffer and cells were pre-extracted with 0.5% Triton X-100 in CSK buffer for 5 minutes on ice. Cells were then fixed in 2% PFA for 15 minutes. (For PCNA labeling, cells were washed twice with PBS then incubated with ice cold methanol on ice for 5 minutes). Cells were washed twice with 1X PBS (1x PBS+1% NGS thrice if antibody labeling was done) for 5 minutes. If EdU was incubated Click-It solution made in 1X PBS containing final concentration 100 mM Na-ascorbate, 10 mM $CuSO_4$, and 1 μM 488-azide was incubated on the coverslip for 1 hour in a covered humidified chamber at RT. If Halo ligand was incubated post fix: 400 nM Halo Ligand JFX 585 in 0.5% BSA in 1X PBS for 1 hour in a covered humidified chamber. For antibody labeling (Mouse-anti-PCNA 1:400; Rabbit-anti-MCM3 1:300) primary antibodies were incubated on the coverslip for 1 hour in a covered humidified chamber at RT. Cells were washed thrice with 1X PBS (1x PBS+1% NGS if antibody labeling was done) for 5 minutes. Secondary antibodies (Goat-anti-Mouse Alexa Fluor 647; Donkey-anti-Rabbit AF 546; Goat-anti-Mouse AF 488; Donkey-anti-Rabbit AF 488 1:1000) were incubated on the coverslip for 1 hour in a covered humidified chamber at RT. Cells were washed twice with 1X PBS and mounted on slide with Vectasheild containing DAPI.

Primary antibodies for IF:

PCNA 1:400 (Santa Cruz CAT# sc-56); MCM3 1:300 (gift from Bruce Stillman CAT# 738); Goat-anti-Mouse Alexa Fluor 647 1:1000 (ThermoFisher CAT# A-21235); Donkey-anti-Rabbit AF 546 1:1000 (ThermoFisher CAT# A-10040); Goat-anti-Mouse AF 488 1:1000 (ThermoFisher CAT# A-11001); Donkey-anti-Rabbit AF 488 1:1000 (ThermoFisher CAT# A-21206)

**Western blot:**

Isolation of Halo-tagged proteins and interactors was performed by immunoprecipitation using the ChromoTek Halo-Trap Magnetic Agarose Kit (Proteintech) according to manufacturer's instruction.

For whole cell lysate cells were grown to 80% confluency then lysed in RIPA buffer supplemented with protease and phosphatase inhibitors. Samples were analyzed with standard Western Blotting protocol.

Primary antibodies for WB:

Actin 1:500 (Santa Cruz CAT# sc-47778); MCM2 1:2000 (gift from Bruce Stillman CAT# 732); panMCM 1:400 (gift from Bruce Stillman CAT# 730); And1 1:1000 (Bethyl A1301-141A-T); Phospho-Chk1 (Ser345) 1:300 (Cell Signaling Technology CAT# 2348); Chk1 1:500 (Cell Signaling Technology CAT# 2345); Orc2 1:500 (pAb 205-6); Rad21 1:500 (Cell Signaling Technology CAT #4321); αTubulin: 1:3000 (Sigma CAT# T5168)

**Flow Cytometry:**

Cells were collected after treatment and fixed with 90% ethanol and incubated at 4°C overnight. Cells are resuspended into PBS 1% NGS and incubated with 10mg/ml RNAse A and 24 µg/ml propidium iodide stain and incubated at 37°C for 1 hour with shaking. Immediately prior to flow analysis, cells are filtered through 5mL Polystyrene Round Bottom Tube with Cell-Strainer Cap (Corning science CAT# 352235). For each sample, 10,000 cells were analyzed by PI flow cytometry.

### Quantification and statistical analyses

Statistical analyses were performed by two-tailed Student's t test unless indicated differently in the figure legends. Quantifications represented as mean ± standard deviation. Further statistical details of experiments can be found in the figure legends.

### MINFLUX data analysis for MCM imaging in fixed cells

The localizations of the 2D imaging sequence were exported from Imspector as a .npy file. A custom python analysis software was used to filter localizations, correct for drift and identify individual MCM molecules as well as further analysis. The filters for the localizations are inspired from literature[56].

For higher sampling of molecules, the third and fourth iterations of the MINFLUX sequence were used.

The center-frequency-ratio (cfr) describes the ratio between emission frequency of the central position of the MINFLUX sequence over the average emission frequency at the outer positions. The cfr can be seen as a quality criterion of the localization process. A 0.8 cutoff for the cfr was previously reported and utilized in this work.

The effective photon frequency at offset (EFO), describes the emission frequency of the fluorophore. The peak of the EFO population was determined to be around 50,000. To remove potential localization of multiple emitters, a cutoff of 80,000 was chosen. During the project, a hardware upgrade increased the laser power, thus a higher cutoff of 90,000 was chosen for these data sets.

To mitigate the impact of sample and beam drift, we utilized the beam line corrected localizations ('loc'). Further, a drift correction algorithm previously used for MINFLUX data[56] was applied on the data and we limited the maximal time per region of interest to 2200s. Lastly ROI were inspected manually for signs of drift.

**Identification of individual molecules**

For identification of individual MCM molecules first a DBscan was performed to find clusters of localizations. To separate molecules within the DBscan detected cluster, K-means clustering algorithm was performed on the DBscan clusters. For K-means the number of clusters was automatically decided by calculating the silhouette score. Additionally, a manual correction for over-separated cluster was performed.

**Simulated distributions**

To model and simulate a random distribution, the alpha shape package was used to outline the MCM positions. Then individual molecules were repositioned within the outline according to a uniform random distribution. Importantly, the number of molecules detected per ROI and used in simulation are the same.

To simulate dimers, a Gaussian distribution was used to simulate the distance between two molecules to model the detected dimer distribution in S phase. A minimum distance of 6 nm was required to model the limitations of the MINFLUX detection and analysis pipeline. The number of dimers was chosen such that the dimer enrichment of NN < 30 nm is around 0% for each FOV. For both cases, monomer and dimer simulations, the simulations were performed 100 times.
